# A statistical framework for defining synergistic anticancer drug interactions

**DOI:** 10.1101/2025.09.29.679166

**Authors:** Diogo Dias, John Zobolas, Aleksandr Ianevski, Tero Aittokallio

**Author notes:** Correspondence: Tero Aittokallio, PhD, Institute for Molecular Medicine Finland, FIMM, P.O. Box 20 (Tukholmankatu 8), FI-00014 University of Helsinki, Helsinki, Finland.

## Abstract

Synergistic drug combinations have the potential to delay drug resistance and improve clinical outcomes in patients with advanced cancers. However, current cell-based screens lack robust statistical assessment to identify significant synergistic interactions for downstream experimental or clinical validation. Leveraging a large-scale dataset that systematically evaluated more than 2,000 drug combinations across 125 pan-cancer cell lines, we established reference null distributions separately for various synergy metrics and cancer types. These data-driven reference distributions enable estimation of empirical *p*-values to assess the significance of observed drug combination effects, thereby standardizing synergy detection in future studies. The statistical evaluation confirmed key synergistic combinations and uncovered novel combination effects that met stringent statistical criteria, yet were overlooked in the original analyses. We revealed cell context-specific drug combination effects across tissue types and inherent differences in statistical behavior of the synergy metrics. To demonstrate the general applicability of our approach to smaller-scale studies, we applied it to evaluate the significance of combination effects in increasing smaller subsamples of an independent dataset. We provide a fast and statistically rigorous approach to detecting synergistic drug interactions in new combinatorial screens, thereby supporting more standardized drug combination discovery.

## Introduction

Drug resistance in cancer renders most monotherapies eventually ineffective, necessitating alternative treatment regimens for relapsed or refractory patients, including synergistic drug combinations, that can overcome resistance mechanisms and improve clinical outcomes [1,2]. Synergistic combination therapies offer several clinical benefits, including enhanced efficacy, reduced likelihood of resistance, and lower toxicities through dose reduction [3].

High-throughput drug combination screening in disease models (cell lines, patient-derived cells, and organoids) has become a standard approach for identifying synergistic drug interactions effects in complex diseases [4-6]. Several anticancer screening studies have evaluated thousands of compound combinations in cancer cell lines and patient-derived cells to assess their co-inhibition effect on cell viability across increasing dose ranges [7-11]. However, even with high-throughput screening (HTS) automation, systematic evaluation of thousands of drug pairs across multiple cellular backgrounds with sufficient replicates becomes rapidly infeasible [8,11]. Furthermore, the scarcity of patient-derived cells further limits the number of combinations that can be evaluated in personalized medicine studies [12]. This results in moderate-sized datasets, where selected drug combinations are tested only in few cell models and doses, often without replicates [7,9,11-14]. Therefore, the current challenge in combinatorial therapeutic discovery is not generating phenotypic drug interaction data, but distinguishing genuine drug combination synergy from experimental noise.

To quantify synergistic effect of the tested combinations, researchers typically rely on established synergy models and software tools, such as SynergyFinder [15] or Combenefit [16], which implement widely used synergy metrics, including Bliss independence [17], Loewe additivity [18], highest single-agent (HSA) [19,20], and zero interaction potency (ZIP) [21]. Despite decades of phenotypic drug combination testing, no standardized approach exists for defining drug combination synergy. Instead, researchers use various statistical and non-statistical approaches [7,11,22-25], which reduces consistency between screening studies, and delays the discovery of truly synergistic drug combinations for therapeutic applications [26]. Arbitrary cutoffs, such as excess over Bliss > 10, can lead to different detection levels between studies [19,23]. These effect-size measures alone cannot determine whether the observed synergy levels are due to statistically significant interactions or random experimental fluctuations. While bootstrapping procedures can provide confidence intervals and empirical *p*-values [27], they rely heavily on the representativeness of the data, often producing biased estimates in small-scale datasets.

These challenges stem from a fundamental gap: researchers lack statistical benchmarks to distinguish true biological synergy effects from random experimental variation. Establishing such benchmarks requires comprehensive reference data from unbiased drug combinations – essentially, a reference null distribution that captures the background variability of synergy metrics under independent drug action. While generating such reference distributions exceeds the resources of most academic laboratories, the Sanger Institute systematically evaluated 2,025 combinations in 51 breast, 45 colorectal, and 29 pancreatic cancer cell lines [13]. These combinations were selected for testing without prior knowledge of potential synergy or antagonism, hence providing a sufficiently large, near-random set of combinations for unbiased statistical evaluation of observed synergy in other studies.

We developed a computational pipeline to leverage these unique data from the Sanger Institute to establish a statistically rigorous framework for defining drug combination synergy. Our approach predicts full dose-combination matrices from sparse experimental data, computes synergy scores using established synergy metrics, and generates tissue-specific reference distributions that enable empirical *p*-value estimation for any observed synergy metric. This approach transforms arbitrarily chosen effect-size thresholds into objective statistical assessments, providing publicly available reference distributions that allow researchers to calculate empirical *p*-values for their own combination experiments. Our framework addresses a fundamental gap in combination therapeutics by establishing the statistical foundation needed to accelerate reliable synergistic combination discovery, biological validation and clinical translation.

## Results

### Reference null distributions for statistical evaluation of drug interaction effects

To overcome the experimental and computational challenges of identifying statistically significant synergistic drug combinations, we developed a computational pipeline that integrates machine learning-based dose-combination effect prediction, synergy quantification, and statistical evaluation through reference null hypothesis distributions (**Fig. 1**). The approach is scalable and statistically robust across diverse drug classes, targets, and tissue contexts.

**Figure 1.**
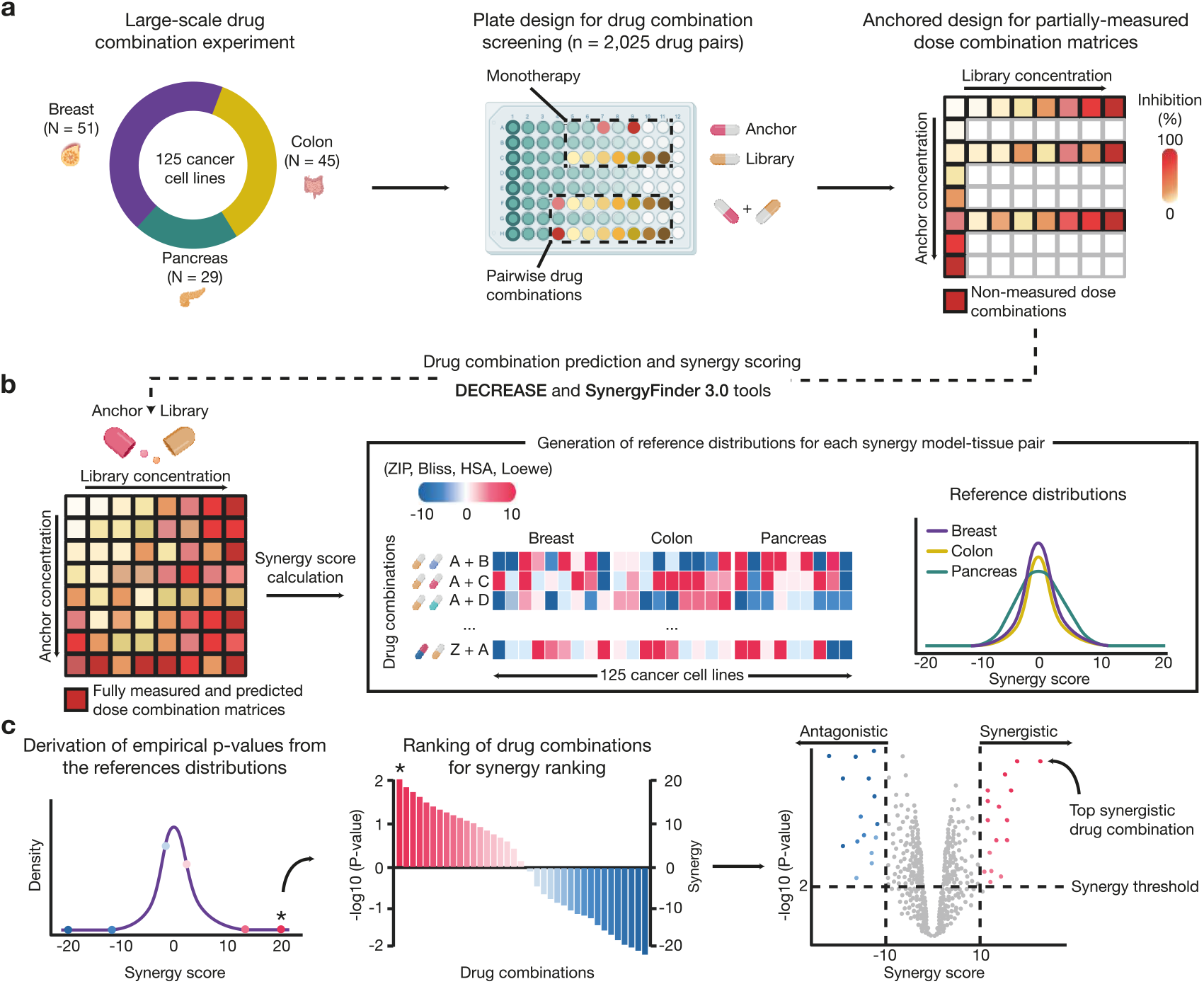
A computational framework for the identification of statistically significant synergistic and antagonistic anticancer drug combinations. (**a**) The Jaaks et al. 2 x 7 “anchored” dose-response drug combination design was used to evaluate *n*=2,025 drug pairs in *N*=125 cancer cell lines, resulting in partially-measured dose-combination matrices. (**b**) *Left:* Based on this large-scale dataset, the DECREASE machine learning model was used to predict the remaining, non-measured combination responses, enabling the subsequent computation of synergy metrics from complete 7 x 7 dose-response matrices using the SynergyFinder 3.0 web-tool. *Right*: reference distributions for the selected synergy metrics were generated across different tissues to derive empirical *p*-value, providing a measure of statistical significance of observed combination effects in new studies. (**c**) In applications of the reference distributions, the newly tested drug combinations are ranked based on their synergy scores (effect size) and empirical *p*-values (statistical significance), with the top-ranked combinations identified as either synergistic or antagonistic hits.

We applied the pipeline to the combination data described by Jaaks et al. [13], which tested 2,025 drug pairs across 125 cancer cell lines as part of the Genomics of Drug Sensitivity in Cancer (GDSC) project [28] (**Fig. 1a**). These high‐throughput screens employed a 2 x 7 anchored dose design with partially measured dose-combination responses. To address the incompleteness of these matrices, we applied our DECREASE machine learning (ML) model [29] to predict complete 7 × 7 dose-combination matrices, after which widely-used synergy metrics (ZIP, Bliss, HSA, and Loewe) were computed for each drug combination and cell line using the SynergyFinder 3.0 web-tool [15] (**Fig. 1b**). A sufficiently large set of multi-dose combination measurements is essential for reliable modelling and quantification of true synergistic and antagonistic effects in tissue-specific combination experiments.

Quality control analyses indicated consistently high assay performance across the screened plates (**Supplementary Fig. 1**; **Supplementary File 1**). Distributions of anchor, library, and combination viability measurements were concordant with assay plate control behavior reported by Jaaks et al. [13], with colorectal cancer plates exhibiting systematically lower viability values and breast cancer plates showing higher viability across conditions (**Supplementary Fig. 1a**). Plate-level quality metrics further demonstrated robust assay performance, with only 162 of 3,106 plates (5%) showing Z′ values below the threshold of 0.5, and none falling below 0.42 (**Supplementary Fig. 1b**). Strictly standardized mean difference (SSMD) values were uniformly elevated across all plates, with no SSMD below 4, indicating excellent separation between positive and negative control wells (**Supplementary Fig. 1c**). This confirms high quality data for the reference distribution modelling. To validate the DECREASE-based dose-combination response predictions, we evaluated the ML predictions using an independent external dataset of fully measured 7 x 7 dose-combination matrices from the pan-cancer study by Bashi et al. [14]. Response matrices were predicted from anchored 2 x 7 designs, mimicking the Jaaks et al. experimental design, and they yielded synergy scores that closely resembled those computed from the fully measured data (overall Spearman correlation *r* = 0.787), with consistent agreement across breast (*r* = 0.775), colorectal (*r* = 0.763), and pancreatic cancer (*r* = 0.837) cell lines (**Supplementary Fig. 3a**). Ranking of the top-synergistic drug combinations was consistently preserved across the tissues (**Supplementary Fig. 3b**), and distributional comparisons showed minimal shifts between predicted and measured ZIP synergy score distributions (**Supplementary Fig. 3c**).

The predicted synergy scores for 2,025 drug combinations across 125 cancer cell lines were next used to establish reference (null) distributions for each tissue type and synergy metric (**Fig. 1b**, right panel). These reference distributions serve as null hypothesis models, capturing the “random” background variability in the tissue-specific combination measurements under the assumption that most of the interactions are additive (i.e., no significant interaction effect). This assumption is supported by the large-scale combination screens showing that synergy is a rare event, occurring in ∼5% of randomly tested drug pairs [6, 10, 14]. Based on these reference distributions, empirical *p*-values can be derived for an observed synergy level (upper tail for synergy, lower tail for antagonism; see **Methods** and **Eq. 1**), providing a measure of statistical significance in any combination screening assay and a given synergy score. The empirical significance testing was well calibrated across tissues, as observed significance rates closely followed the nominal thresholds and the empirical *p-*values showed approximately uniform behavior under the null hypothesis (**Supplementary Fig. 2a** and **b;** see **Methods** and **Eq. 2**). As expected, applying Benjamini-Hochberg false discovery rate (FDR) correction [34] substantially reduced the number of synergistic discoveries compared with nominal significance threshold (**Supplementary Fig. 2c**), underscoring the importance of multiple-testing control when testing a large number of combinations in a given tissue or cell line. This statistical evaluation ensures that only combinations with sufficient statistical evidence of synergy (or antagonism) are prioritized in the follow-up studies, reducing false positives and thereby decreasing the time and costs of the downstream analyses.

Based on the empirical *p*-values (statistical significance) and the observed synergy scores (effect sizes), each drug combination in an experiment can be ranked within its respective tissue context (**Fig. 1c**). Combinations with small *p*-values and high synergy scores are classified as the top synergistic hits, whereas those with significant negative scores are identified as the top antagonistic hits. Integrating statistical significance with effect size prioritizes the most reliable combinations for further mechanistic validation, analogous to differential gene expression analyses that jointly consider both fold changes and adjusted significance levels (volcano plots; **Fig. 1c**, right panel). Such dual-ranking approach facilitates the discovery of significant drug combinations that are likely to exhibit pre-clinically meaningful therapeutic effects when co-administered together in a specific cellular context. Overall, this computational-statistical pipeline establishes tissue-specific reference null distributions that enable statistically principled assessment of synergistic and antagonistic drug combination effects in cell-based viability assays, without requiring additional large-scale measurements across multiple cell lines or concentrations. This pipeline provides a scalable foundation for reference-based statistical evaluation of drug interaction effects in high-throughput drug combination screening data.

### Tissue-specific landscapes of synergistic and antagonistic combination effects

We next examined the distribution of synergy scores and their statistical significance using tissue-specific reference distributions derived from 125 cancer cell lines and 2,025 drug pairs in the Jaaks et al. dataset (**Fig. 2**). This systematic evaluation enabled the identification of novel synergistic and antagonistic interactions by completing full dose-combination matrices through ML-based response predictions. Statistical significance was assessed using empirical *p*-values with FDR control applied within each cell line, and the complete set of adjusted results for all tested drug combinations is provided in **Supplementary File 2**.

**Figure 2.**
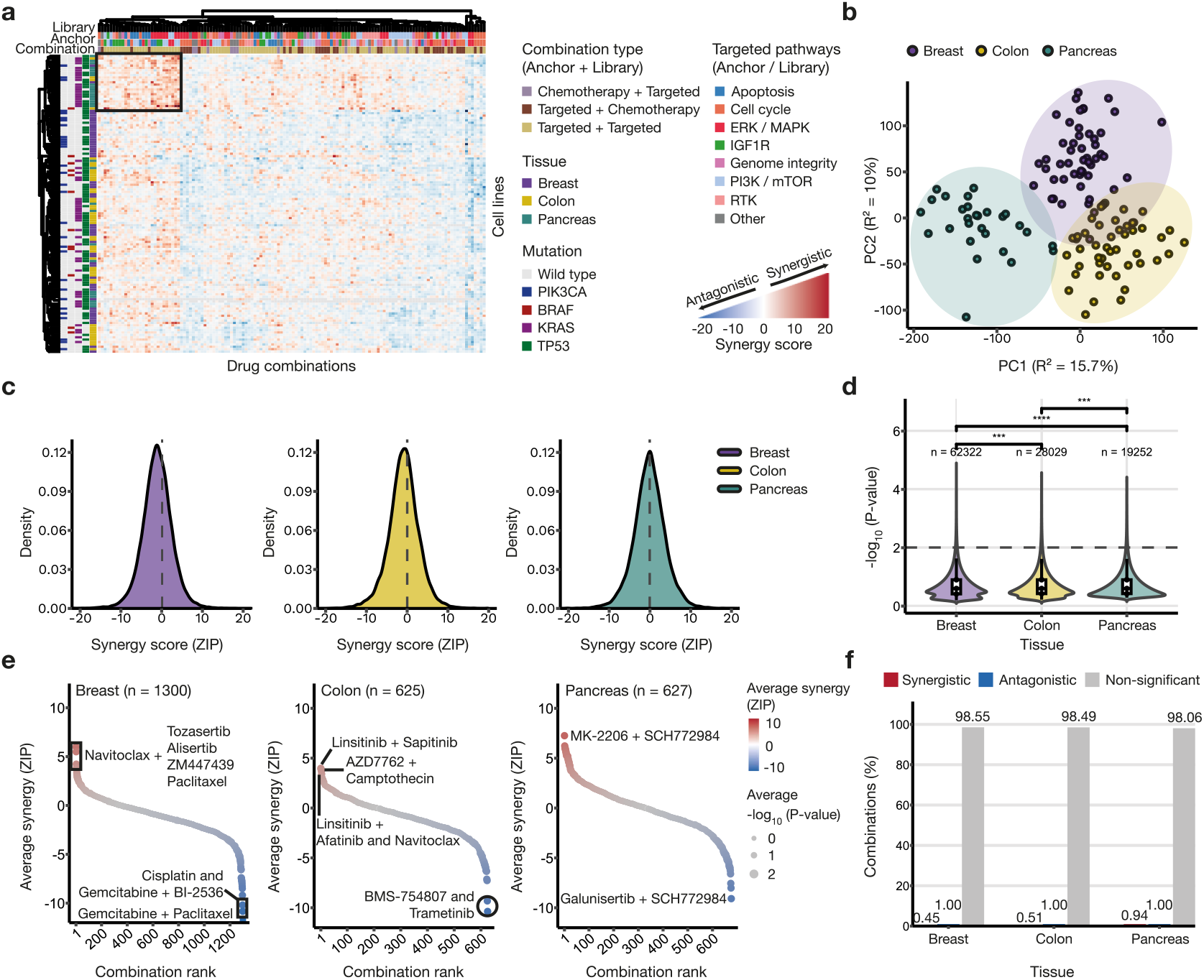
Distribution of synergy scores across breast, colon, and pancreatic cancer cell lines after completing the full drug combination matrices. (**a**) Heatmap of ZIP synergy scores for 125 cell lines and 2,025 drug pairs from the Jaaks et al. dataset, clustered by hierarchical clustering (complete linkage) using a custom Minkowski distance function. For visualization and cross-tissue comparison, drug combinations with >20% missing values were excluded to ensure comparability and reduce bias from incomplete measurements. Only drug pairs for which both anchor and library pathway annotations were available in each tissue were used in the analyses (*N* = 125, *n* = 132). See **Supplementary Fig. 4** for full tissue-specific heatmaps. Colors indicate ZIP scores, with annotations for tissue of origin, key mutations, drug categories, and biological target pathways. (**b**) Principal component analysis (PCA) of the ZIP scores across all drug pairs and cell lines (the colored points). (**c**) Reference (null hypothesis) distributions of the ZIP scores across the three cancer tissues. The dotted horizontal line dashed marks zero ZIP synergy, corresponding to the additive baseline against which synergy is evaluated (synergy and antagonism). (**d**) Violin distributions of log-transformed *p*-values across the tissue types. The central dot indicates the median and the bars correspond to the interquartile range (IQR). The dotted horizontal line marks the selected significance cutoff of *p* ≤ 0.01. Statistical significance was assessed using Wilcoxon rank-sum tests. **\*\*\****p* < 0.001, \*\*\*\**p* < 0.0001. (**e**) Ranking of drug combinations by their average ZIP synergy across cell lines within each cancer type. Each point corresponds to a unique drug pair, with point size indicating the average *p*-value. Top-significant combinations are labelled (both synergists and antagonists). (**f**) Proportion of synergistic, antagonistic, and non-significant drug combinations based on ZIP synergy scores and empirical *p*-values, where synergy corresponds to ZIP ≥ 10 (*p* ≤ 0.01) and antagonism to ZIP ≤ –10 (*p* ≤ 0.01).

We first visualized the landscape of ZIP synergy scores across drug combinations and cell lines in breast (*N* = 51), colorectal (*N* = 45), and pancreatic (*N* = 29) cancers. The analysis revealed distinct patterns of synergy and antagonism stratified by drug combinations and cell line characteristics, including tissue of origin, point mutations, and biological pathways (**Fig. 2a**; **Supplementary File 3**). For example, a distinct synergistic cluster comprising predominantly pancreatic cancer cell lines (black box) was observed, in which mainly molecularly targeted compounds were combined. Although tissue of origin was a major contributor to the observed clustering structure, cell lines did not cluster exclusively by their tissue or key point mutations, highlighting substantial context-specific heterogeneity in drug combination responses both within and across tissues. When analyzed within individual tissues or using alternative synergy metrics, additional differences emerged (**Supplementary Figs. 4** and **5**). Specifically, analyses using the HSA metric showed an inflated synergy landscape, reflecting a known metric bias that positively skews distributions and influences synergy detection. In contrast, analyses with the Loewe metric showed a substantial fraction of combinations for which synergy scores could not be reliably estimated. Correlation analysis across the metrics indicated similar overall behavior between ZIP and Bliss (**Supplementary Fig. 6**; **Supplementary Table 1**). Based on these observations, ZIP was selected as a balanced metric for subsequent analyses, although the statistical framework is broadly applicable to other existing or emerging synergy metrics.

We next examined tissue-specific patterns in drug combination responses using principal component analysis (PCA) of ZIP synergy score profiles. This analysis revealed a clear tissue-specific separation of the cell lines along the major principal components (**Fig. 2b**), indicating that tissue of origin remains a substantial factor of the overall variance in combination synergy. However, this global separation does not imply uniform responses within each tissue, as considerable heterogeneity remains among cell lines of the same cancer type. These findings are consistent with prior reports showing that tissue identity is a dominant, but not exclusive determinant of drug combination response patterns [13,14]. We therefore analyzed the reference null distributions of the ZIP synergy metric separately in each tissue type (**Fig. 2c**; **Supplementary Fig. 7**). The distributions exhibited modest shifts between the tissues, suggesting differences in overall synergy levels by cancer type. The ZIP reference distributions were relatively symmetric and distributed around zero, yet non-normally distributed (**Table 1**), indicating the need for non-parametric statistical assessment of tissue-specific significance. Bliss and ZIP metrics behaved rather similarly, whereas Loewe and HSA produced positively biased reference distributions (**Supplementary Fig. 8**). These reference null distributions form the basis for evaluating the statistical significance of an observed synergy in any combination screen, using a given synergy metric within a specific cancer type. The resulting *p*-value distributions showed an upper-tail enrichment (**Fig. 2d**; *p* ≤ 0.01), consistent with synergy being a rare yet detectable event.

**Table 1.** Distributional characteristics of the ZIP reference null distributions for the three tissue types.

| Tissue | $n$ | Mean | Median | Min | Max | SD | IQR | K-S test |
| --- | --- | --- | --- | --- | --- | --- | --- | --- |
| Breast | 62,322 | -1.130 | -1.110 | -54.0 | 27.5 | 3.73 | [-3.29, 1.08] | $p < 0.001$ |
| Colon | 28,029 | -0.878 | -0.825 | -33.4 | 20.0 | 3.80 | [-3.05, 1.36] | $p < 0.001$ |
| Pancreas | 19,252 | -0.122 | -0.110 | -24.9 | 25.6 | 3.94 | [-2.44, 2.22] | $p < 0.001$ |

$n$ , number of drug combinations evaluated across the cancer cell lines; SD, standard deviation; IQR, interquartile range; KS, Kolmogorov-Smirnov normality test. *Note:* small $p$ -values indicate deviation from the normal distribution, due to large sample size and slightly heavier tails than in normal distribution. The empirical $p$ -value calculation does not require normally distributed synergy scores.

To identify the strongest combination effects within each tissue, we ranked the drug pairs by their average ZIP score across the cell lines (**Fig. 2e**; **Supplementary File 4**). In breast cancer (*n* = 1,300 unique pairs), top synergistic combinations included Navitoclax with Tozasertib, Alisertib, ZM447439 and Paclitaxel. In colon cancer (*n* = 625), the highest-ranking pairs were Linsitinib with Sapitinib, Afatinib or Navitoclax, and AZD7762 with Camptothecin. In pancreatic cancer (*n* = 676), MK‐2206 + SCH772984 emerged as the top synergistic hit, followed by Taselisib + Trametinib, Trametinib + MK‐2206, and Paclitaxel + Navitoclax, with compounds appearing as either anchors or libraries. Similar conclusions were drawn for the antagonistic combinations. Using relatively stringent thresholds for effect size (ZIP ≥ 10 for synergy; ZIP ≤ −10 for antagonism) and empirical significance (*p* ≤ 0.01), we identified a small subset of robust interaction effects (**Fig. 2f**), emphasizing their rarity and tissue context-specificity. The fraction of significant synergistic or antagonistic pairs remained less or equal to 1%, as expected at this *p*-value threshold with the additional requirement for the ZIP cutoff. These cutoffs can be freely adjusted depending on the screening application, allowing researchers to balance sensitivity and specificity. Considering both effect size and empirical significance is therefore essential to distinguish true interactions from additive background.

Together, these analyses delineate tissue-specific landscapes of synergistic and antagonistic drug combination effects and establish reference distributions derived from large-scale combination matrices. These reference models provide a systematic basis for statistically robust evaluation of drug interaction effects and enable the identification of tissue-specific synergistic combinations in future studies using the pre-computed distributions from the Jaaks et al. dataset.

### Identification of significant tissue-specific drug combinations and co-targeted pathways

We next evaluated recurrent combination effects and their associated pathway targets across the three tissue types using the established statistical framework. To identify statistically significant interactions, we applied relatively stringent thresholds for both effect size and empirical significance (synergistic: ZIP ≥ 10, *p* ≤ 0.01; antagonistic: ZIP ≤ –10, *p* ≤ 0.01).

In breast cancer cell lines, the most frequently synergistic anchor drugs were Navitoclax (n = 78; n, number of significant instances), AZD7762 (n = 23), and Linsitinib (n = 20), commonly paired with library compounds such as Gemcitabine and Alisertib (**Fig. 3a**; **Supplementary File 5**). Navitoclax, an inhibitor of anti-apoptotic BCL-2 family proteins, showed particularly strong synergy with Aurora kinase (AURKA) inhibitors, including Tozasertib (n = 11) and Alisertib (n = 10) (**Supplementary Fig. 9**), suggesting coordinated disruption of apoptotic and mitotic control in breast cancer. Importantly, these dominant target pathway associations remained evident after adjustment of empirical *p*-values within each cell line, and the pathway-level enrichments for the key drugs were preserved under FDR control (**Supplementary Fig. 10**). In colorectal cancer cell lines, Linsitinib (n = 19), MK-2206 (n = 14), and AZD7762 (n = 11) emerged as the most frequently synergistic anchors, with enriched combinations involving DNA-damaging agents and inhibitors of the PI3K/mTOR pathway (**Fig. 3a**). Pancreatic cancer displayed a more focused synergy landscape, with Navitoclax again ranking as the top anchor (n = 30), most prominently in combination with the ERK1/ERK2 inhibitor SCH772984 and with Gemcitabine (**Supplementary Fig. 9**). Notably, the HuP-T4 cell line consistently exhibited high synergy, with OSI-027 + Trametinib and SCH772984 + Taselisib among the strongest combinations, highlighting potential benefits of co-targeting ERK/MAPK and PI3K/mTOR signaling in this tissue context. In contrast, the strongest antagonistic interactions were highly tissue-specific and pathway dependent, including Palbociclib with Gemcitabine in breast cancer, suggesting antagonism between cell-cycle inhibition and genotoxic chemotherapy; SB216763 combined with Taselisib or BMS-754807 in colorectal cancer, consistent with suppressive effects from concurrent targeting of WNT signaling alongside IGF1R or PI3K/mTOR pathways; and Galunisertib or LGK974 with SCH772984 in pancreatic cancer, indicating potential suppression through co-inhibition of RTK/WNT and ERK/MAPK signaling.

**Figure 3.**
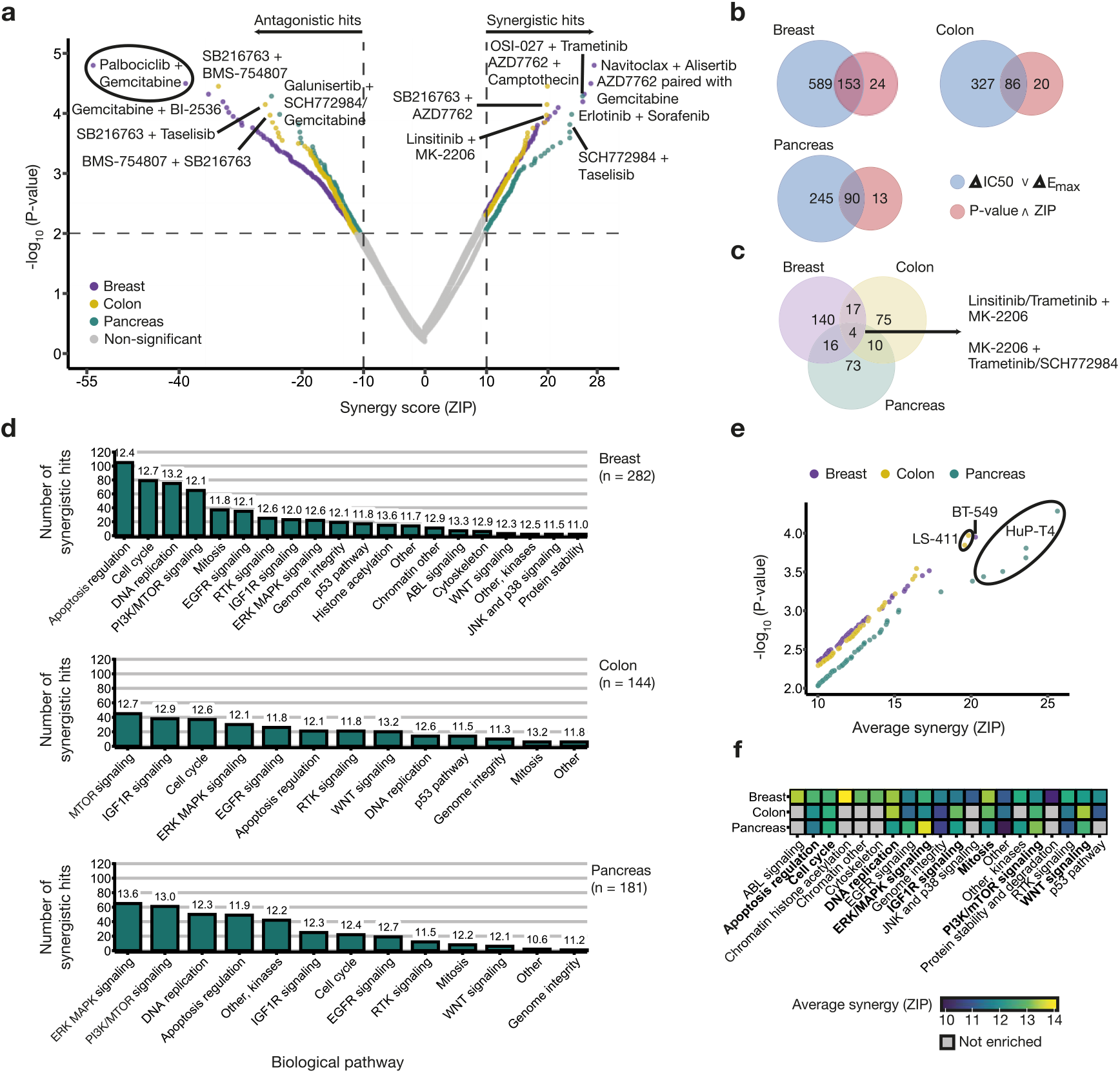
Identification of significant drug combinations and target pathways. (**a**) Volcano plots of ZIP synergy scores and empirical *p*-values for breast, colon, and pancreatic cancer cell lines. Each point corresponds to a cell line-drug pair. The key synergistic and antagonistic drug combinations are highlighted for each tissue, defined as empirical *p* ≤ 0.01 and ZIP ≥ 10 or ZIP ≤ -10, respectively. Dashed lines mark the significance threshold and effect-size cutoff. (**b**) Number of synergistic drug combinations identified based on *p*-values and ZIP scores versus Jaaks et al. detections (△IC50 or △Emax). (**c**) Overlap of highly synergistic drug combinations identified across breast, colon, and pancreatic cancer cell lines (Breast: *n* = 177; Colon: *n* = 106, Pancreas: *n* = 103). Among the tested combinations, only four pairs were shared across the three tissues, each involving targeted agents. (**d**) Number of significant synergistic drug combinations (ZIP ≥ 10, *p* ≤ 0.01) grouped by biological pathway after combining anchor and library drugs. Labels above each bar indicate the average ZIP synergy scores per targeted pathway. Pathways are ordered within each tissue by the number of hits. (**e**) Significant PI3K/mTOR-targeting drug combinations (ZIP ≥ 10, *p* ≤ 0.01) across breast, colon, and pancreatic cancer cell lines. Each point represents a significant drug-pair observation in an individual cell line involving a PI3K/mTOR-targeting agent (Breast *n* = 62, Colon *n* = 40, Pancreas *n* = 56; 89 unique drug pairs across tissues). The three cell lines in which this pathway was most prominently enriched are annotated. (**f**) Top 20 pathways with the highest average synergy. Pathways highlighted in bold represent those commonly observed across multiple cancer types, while grey denotes pathways not enriched or present in the tissue.

To investigate the added value of using empirical *p*-values, we compared the synergistic hits identified by our approach to those reported by Jaaks et al., based on their ΔIC50 and ΔEmax metrics [13] (**Fig. 3b; Supplementary File 6**). Even under the stringent statistical cutoffs, our approach identified 24 additional synergistic combinations in breast cancer, 20 in colon cancer, and 13 in pancreatic cancer cell lines, with a total of 282 synergistic instances for breast, 144 for colon, and 181 for pancreas (**Supplementary File 7**). To further compare synergy definitions between the two approaches, we quantified context-specificity as the number of cancer cell lines in which each drug pair was deemed significant. Across all three tissues, synergies unique to our framework occurred in fewer cell lines than shared synergies, indicating more selective and context-specific discoveries (**Supplementary Fig. 11**). Notably, four common drug pairs demonstrated significant synergy across all three tissues (**Fig. 3c**), each involving the selective pan-Akt inhibitor MK-2206. These combinations occurred either in combination with Linsitinib and Trametinib, or paired with Trametinib or SCH772984, underscoring a pan-cancer synergistic effect from co-inhibition of PI3K/AKT/mTOR and ERK/MAPK signaling. Overall, molecularly targeted combinations dominated the significant synergies across the tissue types, consistent with the observations made by Jaaks et al. [13]. To further characterize tissue-specific pathways underlying statistically significant synergy, we performed pathway-level aggregation of both anchor and library targets within each tissue (**Fig. 3d**; **Supplementary File 8**). In breast cancer, synergistic interactions converged on core cell-intrinsic survival and proliferative programs, with apoptosis regulation, cell cycle progression, and DNA replication representing the most enriched pathways. PI3K/mTOR signaling and mitotic processes further reinforced this proliferation-centric synergy landscape. Colorectal cancer exhibited growth factor-driven synergistic patterns, dominated by PI3K/mTOR and IGF1R signaling, with additional enrichment of ERK/MAPK, EGFR, and cell cycle pathways. Pancreatic cancer revealed a distinct kinase-centered synergy profile, with ERK/MAPK and PI3K/mTOR pathways showing both the highest frequency and the strongest average ZIP synergy.

Notably, the PI3K/mTOR pathway consistently emerged as a central target of synergy across the three cancer types. Highly synergistic combinations within this pathway led to highest synergies, particularly evident for the HuP-T4 pancreatic cell line (**Fig. 3e**). Pathway-level profiling of synergistic combinations identified a conserved set of recurrent processes across tissues, converging on cell survival and proliferative control, with IGF1R and WNT emerging as common drivers (**Fig. 3f**). In addition to the shared programs, tissue-specific signatures were evident; chromatin histone acetylation in breast cancer and ERK/MAPK signaling in pancreatic cancer. The latter was most apparent in HuP-T4, which consistently exhibited strong synergy (**Supplementary Fig. 12**). Antagonistic interactions in breast cancer were uniquely associated with co-targeting of p53 alongside cell-cycle and receptor tyrosine kinase (RTK) inhibition, whereas in colon and pancreatic models the WNT-pathway blockade frequently served as the backbone of antagonistic multi-target regimens (**Supplementary Fig. 13**). Overall, while synergistic responses converge on cell survival and proliferation programs, they are modulated by tissue-specific signaling, reflecting distinct oncogenic wiring and highlighting opportunities for context-specific therapy.

Together, these results demonstrate that the statistical framework enables systematic identification of synergistic and antagonistic combinations that may be overlooked when relying solely on effect size or incomplete experimental measurements. By jointly integrating effect size and empirical significance, the approach supports robust prioritization of tissue-specific drug interactions in combination studies.

### An application of the pre-computed reference distributions to an external dataset

To demonstrate the applicability of tissue-specific reference distributions derived from the Jaaks et al. dataset, we applied the framework to an independent external drug-combination screen from Bashi et al., consisting of 109 fully measured 7 x 7 dose-response matrices [14]. Restricting the analysis to the same cancer types, we further compared empirical significance estimated using the Jaaks reference distributions with bootstrap-based null modeling applied directly to the subsamples of the Bashi dataset (**Fig. 4a**).

**Figure 4.**
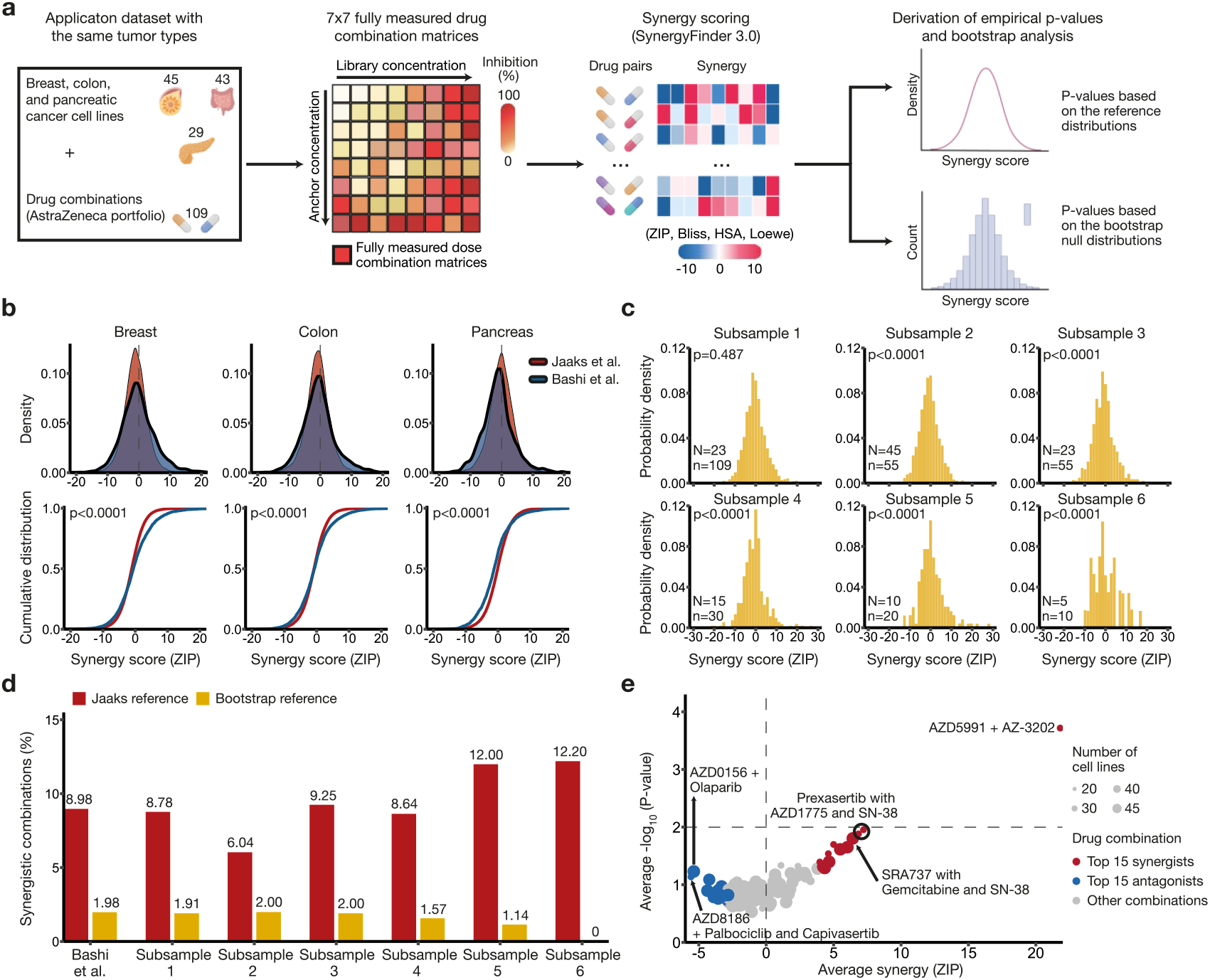
Application of the reference distributions to an external drug combination dataset. (**a**) The Bashi et al. 7 x 7 dose combination dataset was processed to include the same tissues tested as in Jaaks et al, enabling comparison of synergy detection with reference distributions and bootstrap-based analysis. (**b**) Comparison of the ZIP synergy score distributions between the Jaaks et al. and Bashi et al. datasets for each tissue type. *Top*: Probability density distributions of the ZIP synergy scores. *Bottom*: Empirical cumulative distributions of the ZIP scores, highlighting differences in the shape and range of the two datasets. Statistical significance was evaluated with a two-sample Kolmogorov-Smirnov test. (**c**) The bootstrap null distributions obtained by subsampling the full breast cancer dataset of the Bashi et al. (*N* = 45, *n* = 109). Each empirical null distribution was generated from 1,000 bootstrap resamples. Differences between the full and subsampled distributions were assessed using the Kolmogorov–Smirnov test. (**d**) Identification of statistically significant synergies in the Bashi et al. breast cancer cell line dataset based on empirical *p*-values (*p* ≤ 0.01) using two null distributions: the Jaaks et al. reference (red) and the bootstrap reference (gold), each calculated based on 1000 bootstrap resamples per subsample. Bars represent the proportion of statistically significant synergistic combinations in the full Bashi et al. dataset and in six progressively smaller random subsamples, each containing approximately half the number of cell lines and drug combinations than the previous one. (**e**) Volcano plot of average ZIP scores against the average *p*-values in the breast cancer dataset. Each point represents a unique drug combination, with point size proportional to the number of cell lines tested. The dashed vertical line indicates the non-interaction threshold (ZIP = 0), and the dashed horizontal line denotes the significance cutoff of *p* = 0.01.

ZIP synergy scores in the Bashi et al. dataset exhibited substantially heavier tails than their tissue matched distributions from Jaaks et al., with enrichment of both extreme synergistic and antagonistic effects (**Fig. 4b**). This behavior is expected, as the Bashi screen comprised a curated set of prioritized drug combinations, largely drawn from the AstraZeneca portfolio, enriched for compounds with known or suspected antitumor activity, whereas the Jaaks library was assembled without prior knowledge of synergy or antagonism. Consistent with this library design difference, strong enrichment of extreme effects was observed across all tissues (**Supplementary Table 2**), particularly in the tails of the cumulative distribution functions (**Fig. 4b**). As a consequence of this enrichment, the Bashi distributions are not directly suitable as reference nulls without first accounting for the extreme combinations, for example, by identifying and removing significant drug pairs. To further assess cross-study comparability, we focused on drug-cell line perturbations shared between datasets within each tissue, yielding 45 breast, 41 colorectal, and 28 pancreatic cancer cell lines and a total of 189, 64, and 127 shared perturbations, respectively (**Supplementary Fig. 14**).

To evaluate the robustness of null modeling under limited data setting, we subsampled the Bashi breast cancer dataset to approximate the scale of a typical drug combination experiment (**Fig. 4c**). As dataset size decreased, the bootstrap derived null distributions increasingly diverged from those estimated using the full dataset, indicating growing sensitivity to sample composition under reduced data setting. These results indicate that, in small-scale screens, bootstrap resampling can become unstable and may fail to provide a reliable estimate of the underlying background distribution. We next compared how subsampling affected statistical synergy detection under different null assumptions (**Fig. 4d**). Using the Jaaks reference distributions, 8.98% of combinations in the full Bashi dataset were classified as synergistic (*p* ≤ 0.01), with proportions ranging from 6.04% to 12.20% across subsamples. In contrast, the bootstrap-based null distributions showed substantially fewer significant interactions, with detection rates decreasing from 1.91% to zero as sample size decreased. This comparison illustrates how different null modeling strategies can lead to markedly different sensitivity profiles, particularly in smaller combination datasets.

Reflecting on the design of the Bashi compound library, several AstraZeneca compounds appeared among the most synergistic interactions, with the overall landscape skewed toward positive combination effects. Applying our framework, the Mcl-1 inhibitor AZD5991 emerged as a prominent synergistic hit across all tissues, most frequently paired with the Bcl-xL inhibitor AZ-3202 (**Fig. 4e**; **Supplementary Fig. 15**; and **Supplementary File 9**). In addition, multiple DNA damage response targeting combinations were identified in the Bashi dataset, including synergistic interactions between Prexasertib and AZD1775 or SN-38, as well as between SRA737 and Gemcitabine or SN-38. Furthermore, recurrent antagonistic interactions were observed, notably between the ATM inhibitor AZD0156 and the PARP inhibitor Olaparib in breast and colorectal cancer, highlighting pathway-specific efficacy suppression within DDR signaling.

Together, these results demonstrate that large, independently generated tissue-specific reference distributions provide a stable statistical baseline for evaluating external drug combinatorial screens. By contrast, bootstrap-based nulls may lose sensitivity under limited data regimes, underscoring the value of reference-based null modeling strategies for preserving statistical power and robustness.

## Discussion

Drug combination synergy is a relatively simple concept, where two or more drugs when used in combination result in greater than additive effects, compared to using the drugs individually as monotherapies. The complexity originates from the diverse biological mechanisms underlying synergistic interactions, as well as from the variety of metrics used to quantify and interpret combination effects [22,23,30]. To simplify and standardize this process, we introduced a statistical framework to identify robust synergistic and antagonistic drug combinations, building on two established platforms for drug combination prediction and synergy scoring: DECREASE [29] and SynergyFinder [31]. Leveraging partial dose-response matrices from Jaaks et al. [13], the combined use of these two methods enabled the pre-computation of reference distributions from complete 7 x 7 combination matrices across 2,025 drug pairs and 125 cancer cell lines. Regardless of the synergy metric applied, assessing combination effects at multiple doses is critical for defining synergy [23], as single-concentration designs can easily overlook synergistic or antagonistic interactions. Therefore, these reference distributions of close to random synergy scores provide a statistically robust and biologically justified means for evaluating the significance of observed drug interactions through empirical *p*-values. Our data-driven statistical approach enables a standardized definition of synergistic and antagonistic drug pairs in new screens. Importantly, this statistical evaluation enhances the discovery of truly synergistic drug combinations, thereby facilitating a faster and more reliable transition from preclinical to clinical investigation through improved robustness, statistical rigor, and reproducibility.

Even with a relatively stringent cutoff of *p* ≤ 0.01, we identified 24 additional synergistic combinations in breast cancer, 20 in colon cancer, and 13 in pancreatic cancer compared to the original findings from Jaaks et al. [13], underscoring the added value of incorporating empirical *p*-values into synergy detection. Equally important, however, is the removal of false positive findings through statistical synergy evaluation, as spurious detections can lead to wasted resources in downstream validation experiments. The selected cutoffs yielded conservative discoveries, with fewer than 2% of tested combinations classified as significant in the Jaaks dataset using our statistical framework. Importantly, the framework reduces false positives while allowing end users to flexibly adjust significance thresholds according to the intended application (e.g., less stringent cutoffs for exploratory drug discovery versus stricter criteria for confirmation assays). Applying the pre-computed reference distributions derived from Jaaks et al. to the fully measured 7 x 7 pan-cancer dataset of Bashi et al. [14], we further demonstrated the broader applicability of our statistical framework, particularly in smaller-scale experiments where bootstrapping and related resampling procedures become unreliable. Given that drug combination screening is constrained by cost and time, and often limited to only a few dozen combinations tested across small numbers of cell lines and dose levels, the ability to obtain statistically rigorous results under such data-limited conditions provides clear value for drug combination discovery.

To quantify potential synergistic effects, researchers commonly rely on established synergy metrics, such as ZIP, Bliss, HSA, and Loewe. Because these models are based on different assumptions about drug interactions, they may lead to divergent classifications of synergism, additivity, and antagonism. Beyond the selection of the synergy metric itself, researchers must also predefine cutoff thresholds for the synergy detection. Although *ad-hoc* cutoffs, such as ZIP > 10 [32], provide a straightforward means of prioritizing combinations for follow-up investigation, focusing solely on effect size overlooks the statistical significance of the observed synergy, relative to the null hypothesis of no interaction, which quantifies the likelihood that such synergy could arise from random experimental variation alone. Moreover, we demonstrated that reference null distributions can differ substantially across both synergy metrics and tissue types, rendering universal cutoffs inaccurate and contributing to inconsistencies between studies. For example, HSA exhibited a positive bias even for randomly paired drug combinations, raising concerns about its suitability for statistical inference. While the use of empirical *p*-values can partially mitigate such biases (e.g., through non-central reference distributions), we recommend prioritizing other metrics, such as ZIP or Bliss, which display more balanced and statistically well-behaved properties in this context.

The identified combinations highlight the tissue-specific nature of synergistic multi-agent therapies, driven by coordinated effects on cell survival and proliferative control through pathways such as DNA damage response, apoptosis regulation, and PI3K/mTOR signaling. Nevertheless, distinct signaling signatures emerged across tissues, including chromatin acetylation in breast cancer and ERK/MAPK signaling in pancreatic cancer. Pathway-level analyses revealed clear differences in target enrichment across the three cancer types. In breast cancer, synergy was primarily associated with the co-inhibition of cell cycle regulation, chromatin modification, and apoptotic pathways, suggesting that targeting core processes governing proliferation and survival may be particularly effective in this context. In colon cancer, synergistic effects were enriched in cell cycle regulation alongside IGF1R and WNT signaling, pointing to distinct vulnerabilities in these models. In pancreatic cancer, the strongest enrichment arose from combinations targeting the ERK/MAPK and PI3K/mTOR pathways, emphasizing the central role of these signaling axes in shaping therapeutic response.

Collectively, these results indicate that while cell cycle programs represent a shared therapeutic target across tumor types, tissue-specific signaling dependencies modulate optimal strategies for combinatorial treatment. Importantly, these pathway-level enrichments reflect statistical associations based on nominal drug targets, and should therefore be interpreted as hypothesis generating, rather than definitive mechanistic explanations of synergy.

Historically, the development of safe and clinically relevant drug combination strategies has relied largely on empirical testing [3]. High-throughput screening (HTS) in cancer cell line panels, both *in vitro* and *ex vivo*, has enabled more systematic evaluation of large numbers of combinations, whether pre-selected based on clinical observations or guided by biologically informed hypotheses [4,7,8,9]. However, results from large-scale screens consistently show that synergistic drug interactions are relatively rare (approximately 5% of tested drug pairs) and strongly dependent on cellular context and tissue of origin [6,10,13,14]. These observations highlight the need for robust statistical evaluation to reliably identify rare, truly synergistic, and tissue-specific drug pairs. By jointly considering effect size (observed synergy score) and statistical significance (empirical *p*-value), our framework enables application-specific significance thresholds, moving beyond universal cutoffs such as selecting a fixed fraction of the most extreme synergies. This approach provides a scalable strategy for constructing reference distributions from large-scale drug combination datasets and for prioritizing drug pairs that exhibit statistically significant interactions. Once established, these reference distributions can be applied to assess the significance of observed synergy levels in any drug combination experiment, regardless of their scale.

Several open questions remain, including how to extend statistical synergy analysis beyond the three tumor types examined in this work. Since the reference distributions exhibited only modest differences across tissues, one potential strategy would be to pool data from all tissues to construct a proxy reference distribution for other cancer types. As sufficiently large-scale combination screens from additional tissues become available, the same pipeline can then be applied to derive new tissue-specific reference distributions. Previous studies have shown that synergistic interactions are dominated by molecularly targeted drugs [13]. Notably, the current datasets primarily comprise epithelial lineage malignancies, and synergy distributions may differ in cancers with distinct developmental origins, mutational landscapes, or microenvironmental contexts. Careful consideration should therefore be taken to avoid overgeneralizing reference distributions beyond biologically comparable tumor types without recalibration. For simplicity, we pooled all tissue-specific combinations regardless of drug class or target pathway. However, depending on the composition of the drug library in a new study, separate reference distributions could be constructed for specific drug classes or biological pathways. A similar principle applies to dosing, where dose-adjusted reference distributions could be generated to better match the concentration ranges of interest. Alternatively, DECREASE [29] and related machine learning approaches [33] can extrapolate beyond the measured dose matrix to expand the effective concentration window. Finally, in studies where healthy (e.g., non-cancerous) cells are profiled using the same drug combinations, the framework could be extended to infer selective synergy in cancer cells relative to toxicity in healthy cells [12], or to define therapeutic dosage windows that optimally balance efficacy and toxicity.

Given these wide possibilities, this work provides a statistically grounded framework for the standardized identification of truly synergistic drug combinations, supporting their prioritization for downstream validation in patient-derived models. By explicitly accounting for cellular context and empirical significance, our approach addresses key challenges in drug combination discovery, offering a robust foundation for the development of personalized healthcare strategies. This is particularly relevant in cancer research, where inter-patient heterogeneity and context-dependent drug responses limit the effectiveness of *one-size-fits-all* therapeutic approaches.

## Materials and Methods

### Drug combination screening data

Jaaks et al. [13] generated a large-scale drug combination dataset comprising 2,025 drug pairs tested across 125 cancer cell lines (51 breast, 45 colorectal, and 29 pancreatic). In this dataset, drug combinations were screened using a so-called 2 x 7 anchored design, in which only a subset of the full drug-dose combination matrix was experimentally measured (see **Fig. 1a**). Each screening plate included a single replicate of the combination dose-response measurements, five replicates of the anchor drug, and four replicates of the monotherapy (single-agent) treatments. Cell viability response to drug treatments was assessed with CellTiter-Glo 2.0 (CTG 2.0, Promega) assays following the transfer of cells into 1,536-well microplates.

The screens were conducted using the Genomics of Drug Sensitivity in Cancer (GDSC) platform [28], generating thousands of viability measurements per drug combination across all cell lines. **Table 1** summarizes the number of drug combinations evaluated across the cell lines. The number of combinations varied between the tissues, due to the variable number of anchors and libraries tested on each plate, and since some of the combinations were not evaluated in each cell line. This dataset provided a suitable large-scale resource of effectively random combinations for constructing reference distributions of synergy scores for each of the three tissues (breast, colon, and pancreatic cancer).

The GDSC platform and CTG 2.0 assays were also used to generate another large-scale dataset by Bashi et al. [14], consisting of fully-measured drug-dose combination matrices for 109 selected anticancer drug pairs from the AstraZeneca’s oncology library evaluated across 755 pan-cancer cell lines using a full 7 x 7 dose-response matrix design. This dataset was used to apply and test the empirical *p*-values derived for observed synergy scores, using the reference distributions estimated from the Jaaks et al. dataset. Among other tissues, the Bashi et al. cell lines included breast carcinoma (*n* = 45), colorectal carcinoma (*n* = 43), and pancreatic carcinoma (*n* = 29).

### Drug combination data preprocessing

In the Jaaks et al. dataset [13], the 2 x 7 anchored drug combination dose-response data were extracted from each screening plate, along with the corresponding monotherapy measurements for both the low and high anchor doses of the first drug and the seven-point concentration series of the library compound (i.e., the second drug in the pair). To complete the monotherapy profiles for the anchored drug, the missing dose-response data were retrieved from other screening plates in which the same anchored drug had been tested as the library drug in the 2 x 7 combination design. The median of the replicate measurements was used to compute the final viability values for both anchored and library monotherapy treatments. In some cases, certain compounds in this dataset included one or two extra concentrations in the monotherapy screens beyond the high/low anchor concentrations. As a result, some matrices expanded to 8 x 7 or 9 x 7 formats, which were also incorporated into our analysis. For the fully measured 7 x 7 drug combination matrix dataset from Bashi et al [14], the drug-dose response viability data for each combination and cell line were directly extracted from the respective screening plates for synergy computation, without any additional processing or ML predictions.

### Assay quality control and plate-level performance metrics

Assay quality control (QC) was assessed at the plate level using established statistical metrics that quantify signal separation and assay robustness based on control of populations. For each screening plate, QC statistics were computed from the negative control (NC) and positive control (Blank) wells, prior to downstream modeling and statistical analysis, consistent with the original viability quantification [13]. Plate robustness was evaluated using the standardized Z′ factor in high-throughput assays defined as: Z′ = 1 − 3 × (σ_p_ + σ_n_) /|μ_p_ − μ_n_|, where μ_p_ and μ_n_ denote the mean responses of the positive and negative control wells, respectively, and σ_p_ and σ_n_ their corresponding standard deviations. Standard deviations were estimated as σ = √(var(*x*) × (*n* − 1) /*n*), where *x* represents the vector of control measurements and *n* denotes the number of non-missing control wells. To reduce sensitivity to outliers and deviations from normality in control distributions, a robust Z′ factor was additionally computed by replacing means with medians and population standard deviations with median absolute deviations (MAD), yielding robust Z′ = 1 − 3 × (MAD_p_ + MAD_n_) /|median(P) − median(N)|. Signal separation between control wells was further quantified using the strictly standardized mean difference (SSMD), defined as SSMD = (μ_p_ − μ_n_) /√(var(P) + var(N)), where var(P) and var(N) denote the sample variances of the positive and negative control responses, respectively. SSMD provides a scale-independent measure of effect size and directly reflects the degree of separation between control distributions.

### Drug combination prediction and synergy quantification

To complete the partially-measured drug combination matrices in Jaaks et al. dataset [13], the 2 x 7 dose-response combination measurements were used as input for the DECREASE machine learning algorithm; algorithmic details and validation results are provided in the original study [29]. DECREASE successfully predicted the full set of 7 x 7 pairwise drug-dose combination response matrices across all 125 cancer cell lines. The predicted combination matrices were then processed using the SynergyFinder v3.0 [15] to quantify synergy and antagonism using the Bliss, highest single-agent (HSA), Loewe and ZIP models. For Bashi et al. [14], synergy scores were directly computed from the 7 x 7 drug combination matrices without requiring matrix prediction.

### Rank stability of prioritized drug combinations following DECREASE predictions

To evaluate the robustness of drug-pair prioritization under reduced experimental designs, we compared drug combination rankings derived from the DECREASE-predicted ZIP synergy scores with rankings obtained from the experimentally measured scores at the drug pair level using the Bashi *et al*. dataset [14]. For each tissue, ZIP scores were aggregated across cell lines to produce a single score per drug combination, resulting in rankings of 109 combinations per tissue. Agreement between predicted and measured rankings was quantified using Spearman’s rank correlation. Prioritization stability was further assessed using a top-N overlap approach, in which drug combinations were ordered by decreasing mean ZIP score and the overlap between predicted and measured top-ranked hits was calculated for varying N values. All analyses were conducted separately for individual tissues and for pooled data across tissues.

### Calculation of empirical *p*-values from the reference null synergy distributions

Empirical *p*-values were computed using nonparametric, tissue-specific reference distributions of synergy scores derived from the Jaaks et al. dataset [13]. For each tissue type, reference synergy scores were constructed separately for each synergy metric (ZIP, Bliss, HSA, and Loewe) and pooled across all screened drug combinations and cell lines to define tissue-level null distributions. Statistical significance of an observed synergy score was assessed by comparing it to the corresponding tissue specific reference distribution, with the total number of reference scores defining the effective sample size. Directional empirical *p*-values were computed to reflect the direction of the observed effect. For positive synergy scores, significance was evaluated using an upper tail empirical *p*-value, defined as the fraction of reference scores greater than or equal to the observed value. For negative scores, corresponding to antagonistic effects, a lower tail empirical *p*-value was computed analogously. To avoid zero *p*-values arising from finite reference sample sizes, *p*-values of zero were replaced by a minimum value equal to the inverse of the reference sample size. This empirical framework is nonparametric and does not rely on distributional assumptions. Additionally, two-sided empirical *p*-values were computed for supplementary analyses to assess deviations in both directions relative to the reference distribution. The empirical *p*-values were computed as:

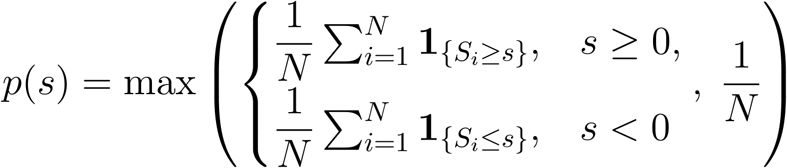

Equation (**1**). Directional empirical *p*-value from tissue-specific reference distributions.

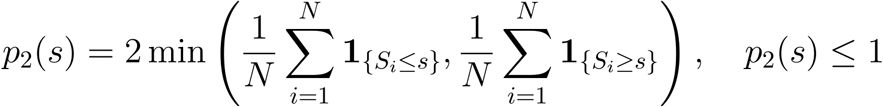

Equation (**2**). Two-sided empirical *p*-value from tissue-specific reference distributions.

where

**N:** the total number of finite reference synergy scores in the tissue-specific null distribution.

**S**_**i**_: the *i*-th reference synergy score in the null distribution for a given tissue (*i* = 1, …, N).

s: the observed synergy score for the evaluated drug combination.

**𝟙{S**_**i**_ **≥ s}:** indicator function equal to 1 if S_i_ ≥ s and 0 otherwise (upper-tail test for synergistic effects).

**𝟙{S**_**i**_ **≤ s}:** indicator function equal to 1 if S_i_ ≤ s and 0 otherwise (lower-tail test for antagonistic effects).

**p(s):** the empirical one-sided p-value computed relative to the reference distribution (upper tail for synergy, lower tail for antagonism).

**p**_**2**_**(s):** the empirical two-sided p-value computed as twice the minimum of the lower- and uppertail probabilities derived from the reference distribution.

### Generation of bootstrap null distributions

To obtain smaller-scale data matrices from the Bashi et al. dataset, which better mimic the sample sizes of a typical drug combination experiment, we randomly subsampled both its drug combinations and cell lines without replacement. More specifically, from the full Bashi et al. dataset **X**, we draw a subsample **X\*** of size *N* x *D* independently with equal probability for any given cell line - drug combination pair. For each subsample **X\***, we then performed bootstrap resampling with replacement to generate 1,000, 2,000, or 5,000 resamples of the same size (*N* x *D)*. To derive the bootstrap null distributions, we primarily used 1,000 bootstrap resamples, as results were highly similar to those obtained with 2,000 or 5,000 resamples. The resulting null distributions were compared with those derived from the full Bashi et al. dataset using the Kolmogorov–Smirnov test. Bootstrap *p*-values were computed the same way as the empirical *p*-values from the Jaaks et al. data reference distributions (**Eq. 1**). The generation of 1,000, 2,000 and 5,000 bootstrap samples required approximately 23 minutes, 47 minutes, and 2 hours per tissue, respectively, when executed on the Puhti high-performance computing at CSC.

## Data and code availability

Jaaks et al. [13] large-scale drug combination dataset can be freely accessed at https://figshare.com/articles/dataset/Original_screen_All_tissues_raw_data_csv_zip/19141916?file=34010816. Bashi et al. [14] drug combination dataset can be freely accessed at https://figshare.com/articles/dataset/Results_recorded_for_each_well_in_7×7_matrix_experiments_for_all_screens/25266697?file=45451363. Analysis codes for the manuscript figures and direct application of the reference distributions is available in the following repository: https://github.com/dias-dio/defining-synergy

## Acknowledgments

We thank the authors of Jaaks et al. and Bashi et al. articles for making their combination data publicly available. The authors thank the CSC – IT Center for Science, Finland, for providing the computational resources. Funding support: AI: Ida Montin Foundation grant. TA: Research Council of Finland (grants 340141, 344698, 367855); the Cancer Society of Finland, the Norwegian Cancer Society (grants 216104 and 273810), Norwegian Health Authority South-East (grants 2020026 and 2023105), the Sigrid Jusélius Foundation, and iCAN – Digital Precision Cancer Medicine Flagship (iCAN-MULTIDRUG). Some graphical elements of Figure 1 were created with the help of the BioRender software.

