## Supplementary Information for "A statistical framework for defining synergistic anticancer drug interactions"

#### Supplementary Figures

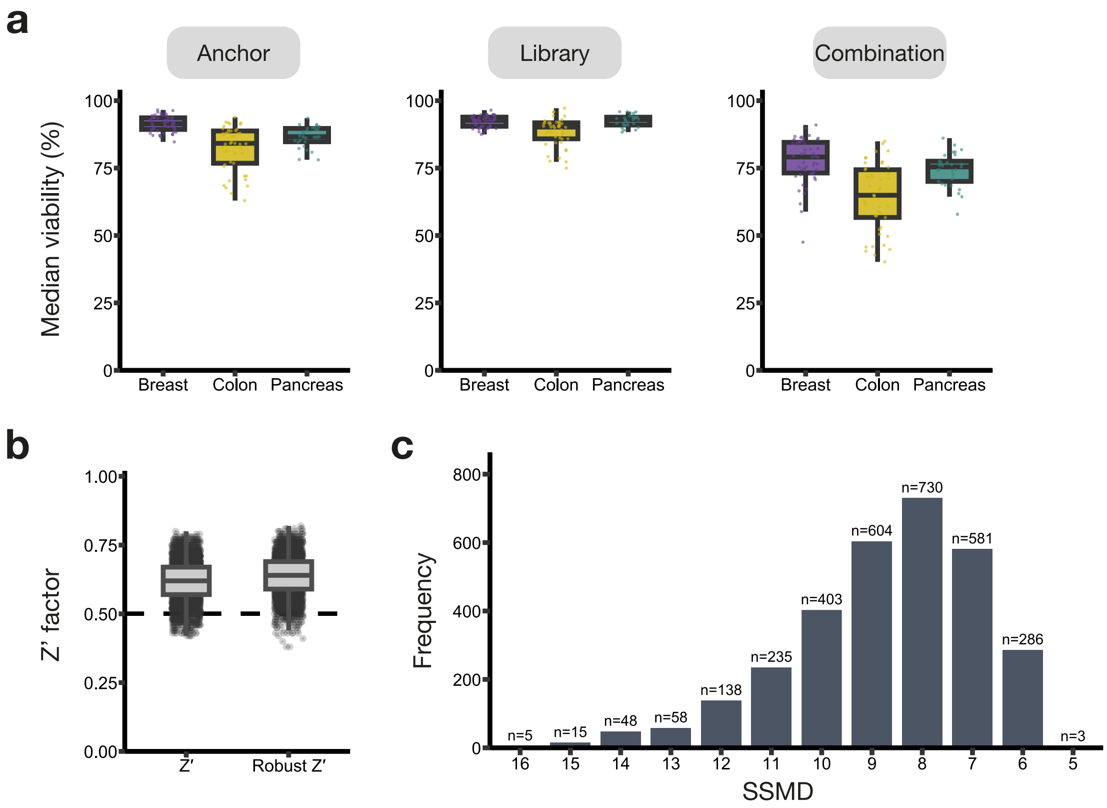


**Supplementary Figure 1. Quality control assessment of viability measurements and assay performance for the derivation of the reference distributions per cancer type.** (**a**) Distribution of anchor, library, and combination viability measurements across tissues. For each tissue, boxplots summarize the median and interquartile range of viability values, with overlaid points representing viabilities. (**b**) Plate-level assay quality assessed using the standard Z′ prime factor and the robust Z′ prime statistic. Each point represents a single screening plate, with paired values shown for the standard and robust Z′ metrics. The dashed horizontal line at Z′ = 0.5 indicates the commonly accepted threshold for excellent assay quality, with values above this line reflecting high assay robustness. (**c**) Frequency distribution of strictly standardized mean difference (SSMD) values across all screened plates. Bars indicate the number of plates with a given SSMD value (rounded to the nearest integer), ordered from highest to lowest frequency. The predominance of high SSMD values highlights consistently strong separation between positive and negative control wells and overall excellent assay performance.

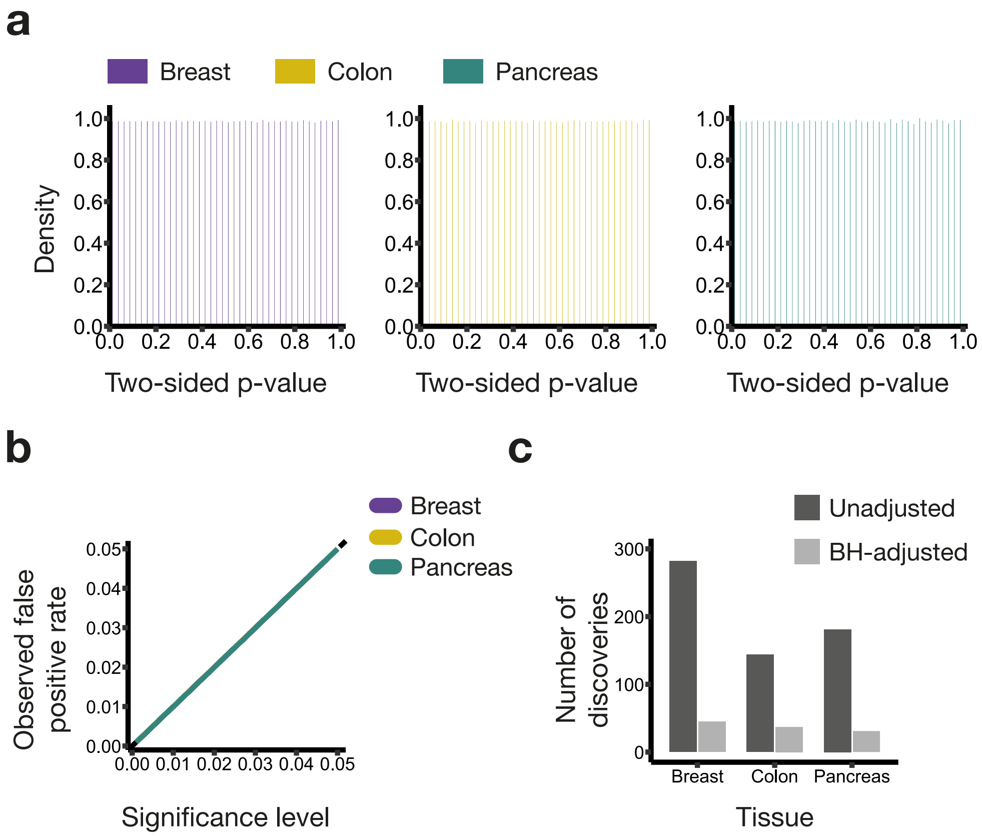


**Supplementary Figure 2. Calibration and significance assessment of drug combination synergy across cancer tissues.** (**a**) Histograms show the distribution of two-sided empirical *p*-values for the ZIP synergy scores in breast, colon, and pancreatic cancer cell lines. *P*-values were computed by percentile ranking of observed ZIP scores against tissue-specific empirical reference distributions, without logarithmic transformation or other corrections. The uniform distributions indicate proper calibration of the empirical hypothesis testing framework. (**b**) Calibration curves relating the nominal significance level to the observed fraction of combinations declared significant based on two-sided empirical *p*-values. The diagonal indicates ideal calibration under the null hypothesis. (**c**) Number of synergistic drug combinations identified using unadjusted nominal *p*-values against those retained post Benjamini-Hochberg false discovery rate (BH-adjusted FDR) correction within each tissue. Synergistic interactions were defined using a ZIP score threshold (ZIP ≥ 10) together with either a nominal significance cutoff ($p \leq 0.01$) or an FDR adjustment ($q \leq 20\%$), illustrating the expected reduction in discovery numbers after the multiple-testing correction.

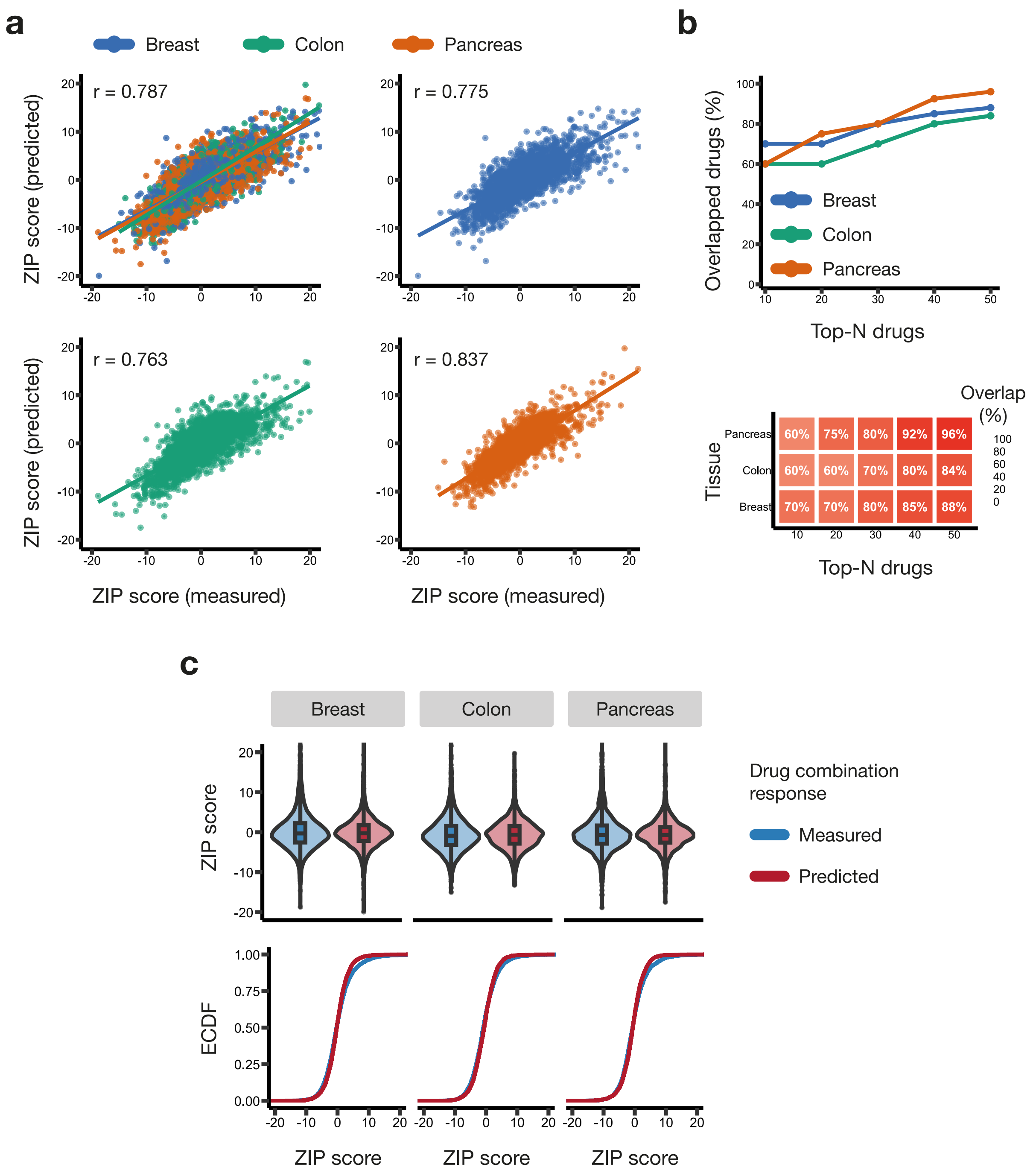


**Supplementary Figure 3. Distributional agreement and rank stability between fully measured and DECREASE-predicted ZIP synergy scores across breast, colon, and pancreatic cancer cell lines.** (**a**) Scatterplots showing the relationship between measured ZIP synergy scores derived from fully measured 7 x 7 drug combination matrices and ZIP scores predicted from reduced 2 x 7 designs using the DECREASE framework, where a subset of dose-response combination measurements was systematically removed from the original 7 x 7 matrices. Each point corresponds to a drug combination-cell line pair (Breast: n = 4,027; Colorectal: n = 4,047; Pancreatic: n = 2,647; total n = 10,721). Across tissues, 109 unique drug combinations were evaluated in 117 cell lines. ZIP synergy scores were computed using SynergyFinder 3.0. Correlations were assessed using Spearman's rank correlation with the *complete.obs* option in R, ensuring that only drug-cell pairs with non-missing values in both datasets were included. (**b**) Rank stability of prioritized drug combinations between predicted and measured synergy scores. Top-N overlap analysis shows the proportion of shared drug combinations between the top-ranked predicted and measured hits as a function of N, evaluated separately for each tissue (*top* plot). The corresponding heatmap (*bottom* plot) summarizes overlap percentages across tissues and N values, highlighting the relatively high preservation of drug-pair prioritization under the reduced 2 x 7 experimental designs. (**c**) *Top*: Violin plots showing the distributions of ZIP synergy scores computed from the fully measured 7 x 7 drug-combination matrices and from 2 x 7 derived predictions for breast, colorectal, and pancreatic carcinomas. Violin width reflects score density, with overlaid boxplots indicating the median and interquartile range. *Bottom*: Empirical cumulative distribution functions (ECDFs) of the measured and predicted ZIP scores for each tissue, illustrating close agreement between the distributions.

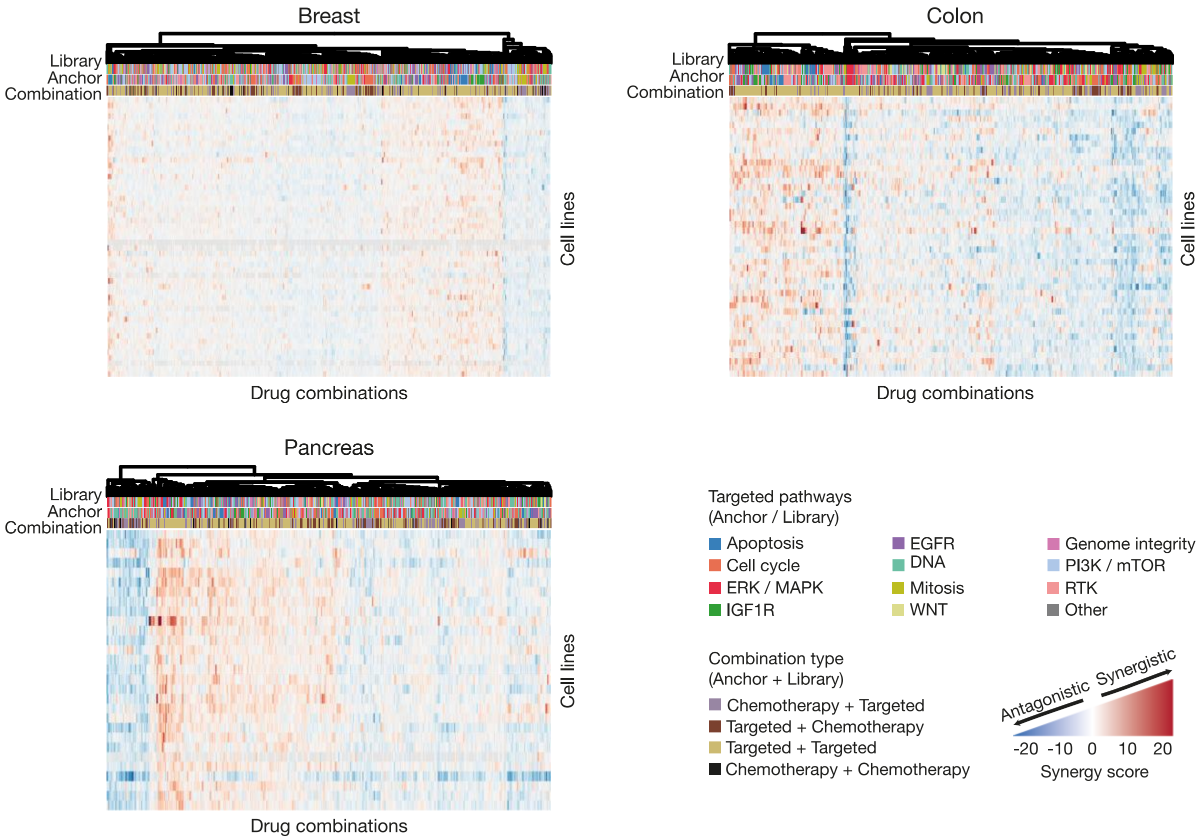


**Supplementary Figure 4. Tissue-specific landscapes of ZIP synergy scores across cancer cell lines.** Heatmaps display ZIP synergy scores across 51 breast cancer cell lines (882 drug combinations), 45 colorectal cancer cell lines (528 combinations), and 29 pancreatic cancer cell lines (506 combinations) from the Jaaks et al. dataset. Drug combinations are clustered along the top dendrogram using hierarchical clustering with complete linkage and a custom Minkowski distance metric.

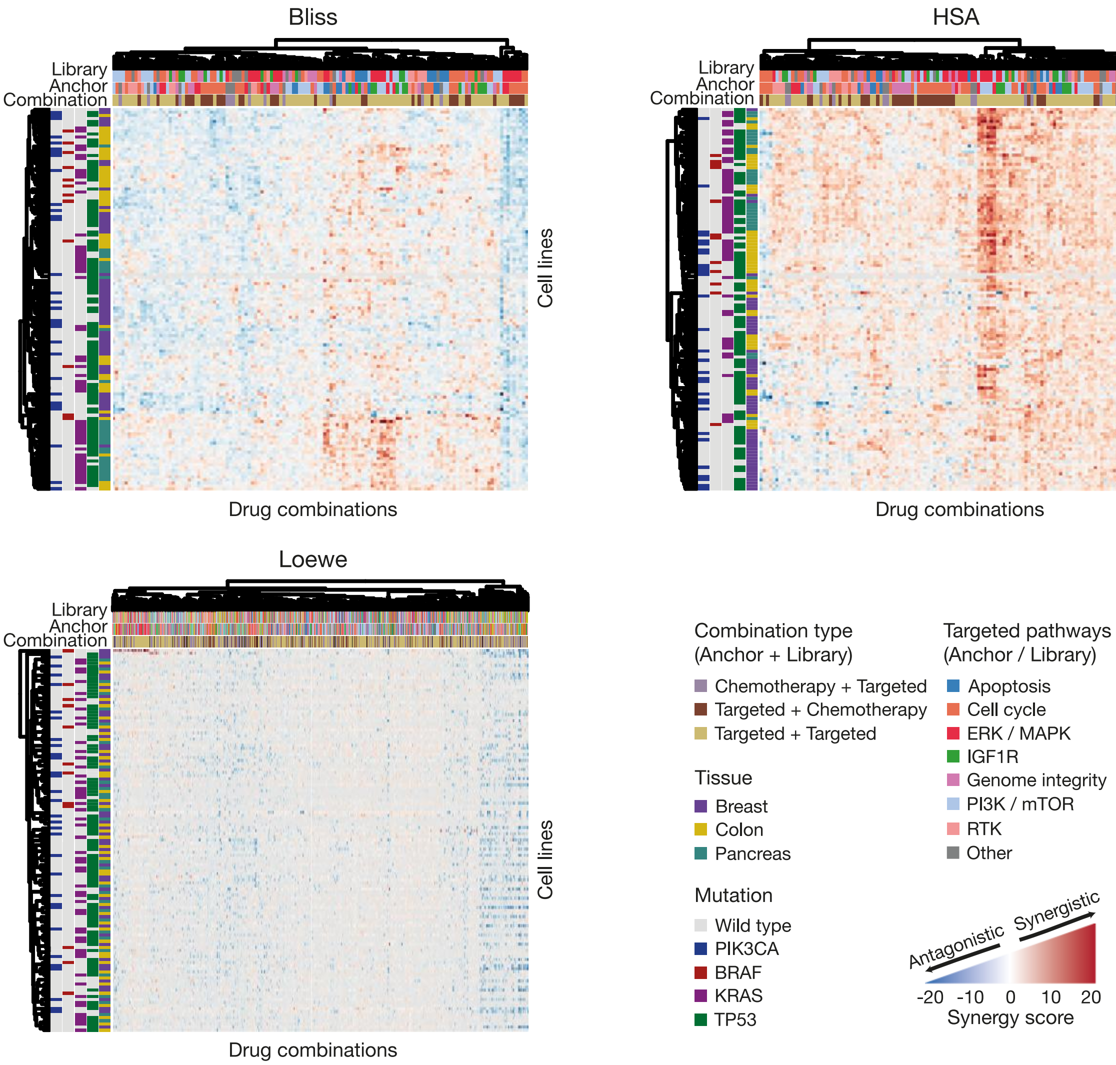


**Supplementary Figure 5. Comparison of Bliss, HSA, and Loewe synergy landscapes across cancer cell lines.** Heatmaps show Bliss, HSA, and Loewe synergy scores across 125 cancer cell lines and drug combinations from the Jaaks et al. dataset. Both drug combinations and cell lines were clustered using hierarchical clustering with complete linkage and a custom Minkowski distance metric. For ZIP and Bliss, drug combinations with more than 20% missing values across cell lines were excluded, whereas for Loewe, combinations with up to 80% missing data were retained. Sample sizes were as follows: Bliss (N = 125 cell lines, n = 132 combinations), HSA (N = 125, n = 132), and Loewe (N = 125, n = 947).

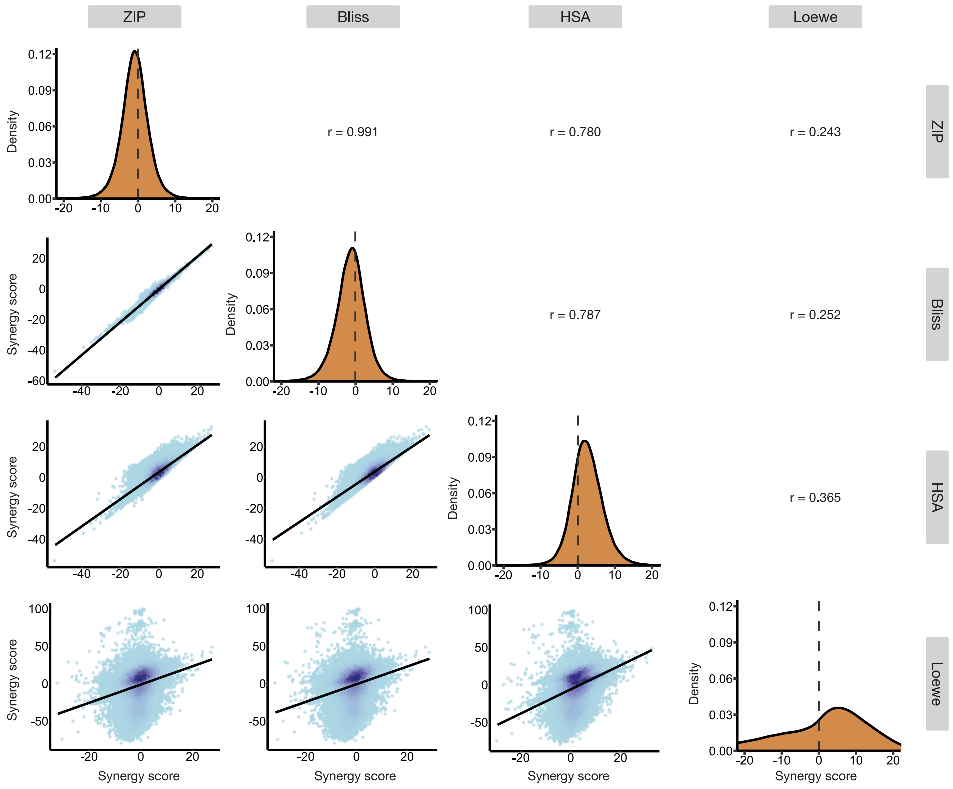


**Supplementary Figure 6. Pairwise correlation of synergy metrics across cancer cell lines.** Scatterplots show correlations between ZIP, Bliss, HSA, and Loewe synergy scores computed across all cancer cell lines. Each point represents a drug combination, with color indicating local point density; darker blue denotes regions of higher density. Point densities were estimated using a two-dimensional kernel density estimator (kde2d function, MASS R package).

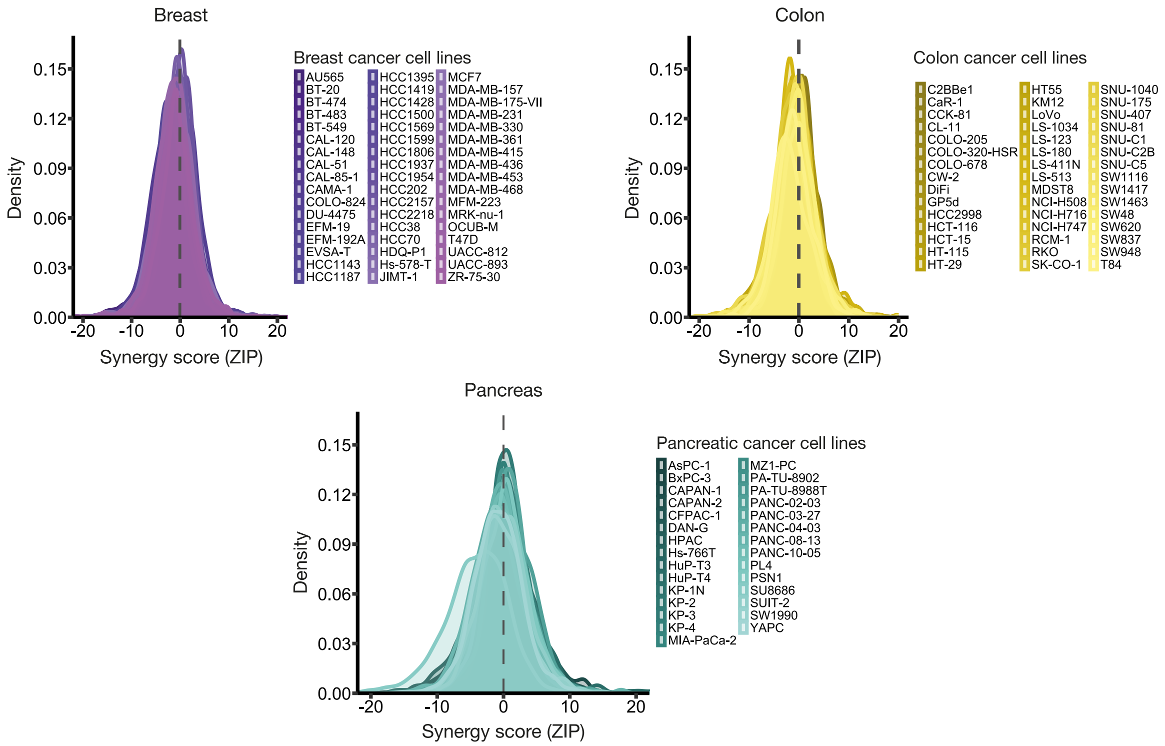


**Supplementary Figure 7. Tissue-specific distributions of ZIP synergy scores.** Density curves show the distribution of ZIP synergy scores across breast, colorectal, and pancreatic cancer cell lines. The dashed vertical line indicates the non-interaction threshold (ZIP = 0). Colors denote tissue types.

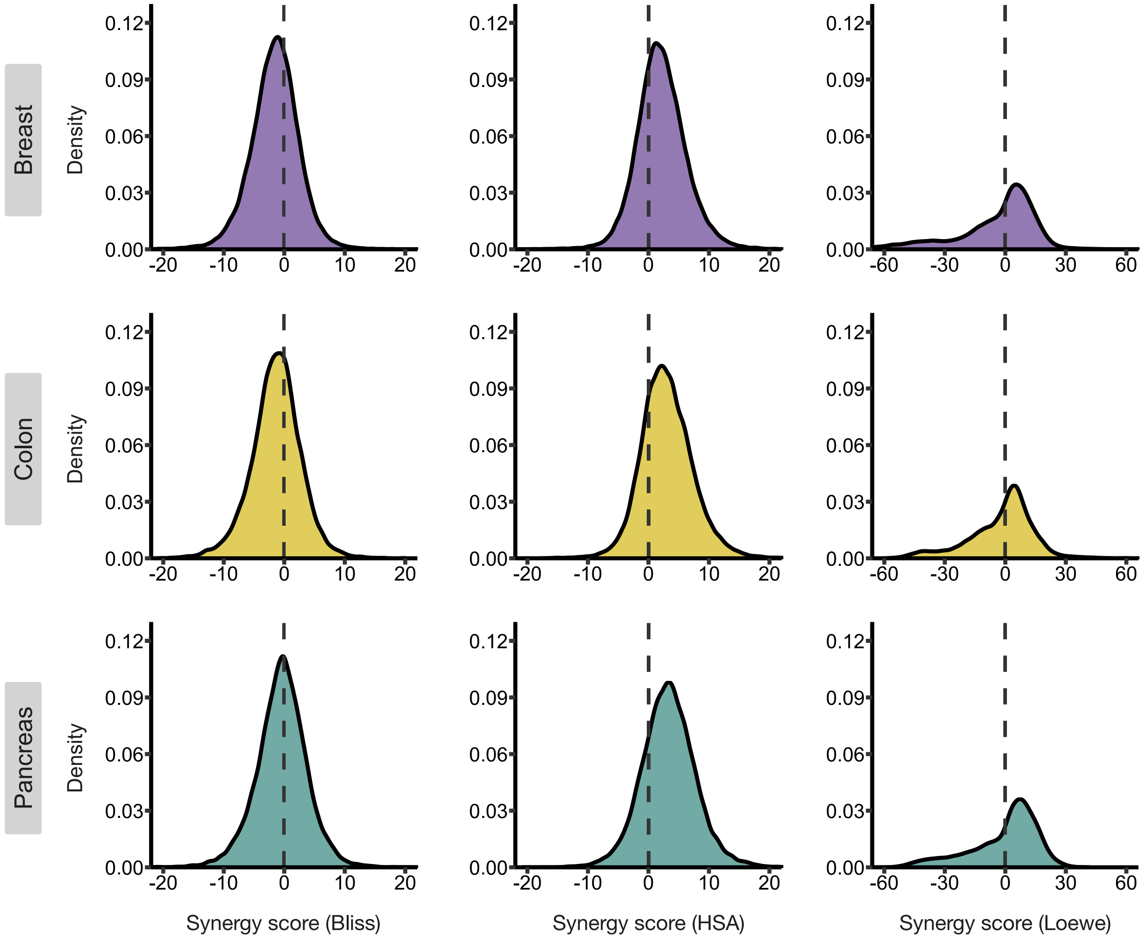


**Supplementary Figure 8. Tissue-specific reference distributions of Bliss, HSA, and Loewe synergy scores.** Density plots show the empirical reference (null hypothesis) distributions of Bliss, HSA, and Loewe synergy scores across breast, colorectal, and pancreatic cancer tissues derived from the Jaaks et al. dataset. Note the different x-axis range for the Loewe distribution, reflecting the smaller number of drug combinations evaluable with this metric. Colors denote tissue types.

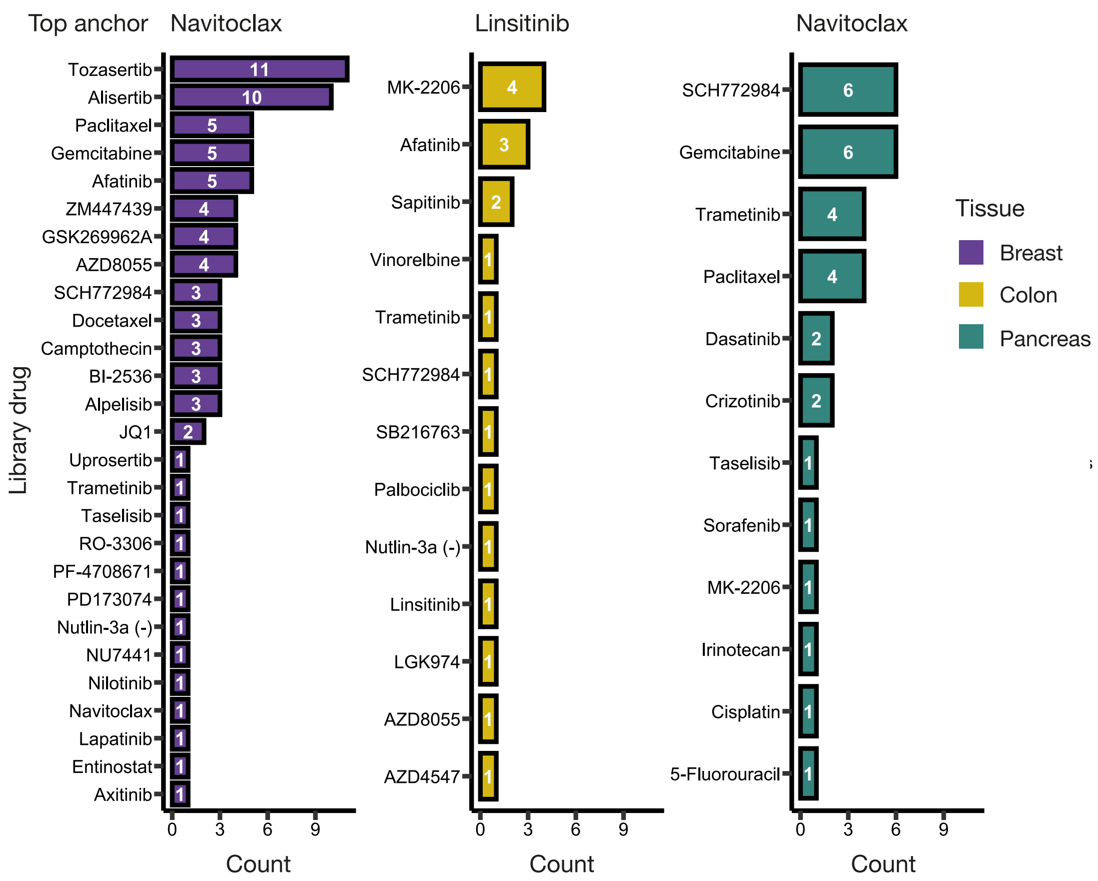


**Supplementary Figure 9. Anchor-level enrichment of synergistic drug combinations across tissues.** Bar plots show the number of distinct library partner compounds associated with each top anchor drug among highly synergistic combinations identified in the Jaaks et al. dataset, stratified by tissue type. This summary highlights anchor compounds that repeatedly form synergistic interactions with multiple partners within a given tissue context.

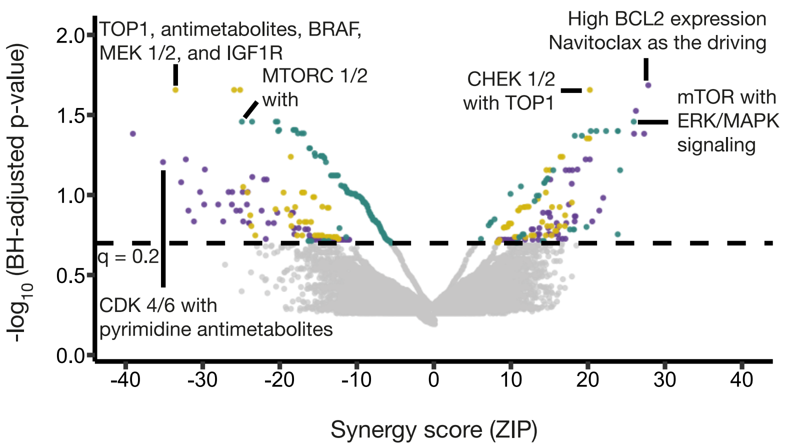


**Supplementary Figure 10. Target pathway prioritization under Benjamini-Hochberg (BH) false discovery rate (FDR)–controlled empirical significance.** Volcano plots summarize drug-combination effects across tissues, with ZIP synergy scores plotted against $-\log_{10}$(BH-adjusted, one-sided empirical *p*-values). Each point represents a cell line-drug pair; non-significant combinations ($q$ > 20%) are shown in grey, while BH-FDR–significant combinations ($q \leq 20\%$) are colored by tissue of origin. For each tissue, the most synergistic and antagonistic combinations are annotated with their corresponding drug targets, highlighting prominent pathway-level interaction patterns preserved under multiple-testing correction.

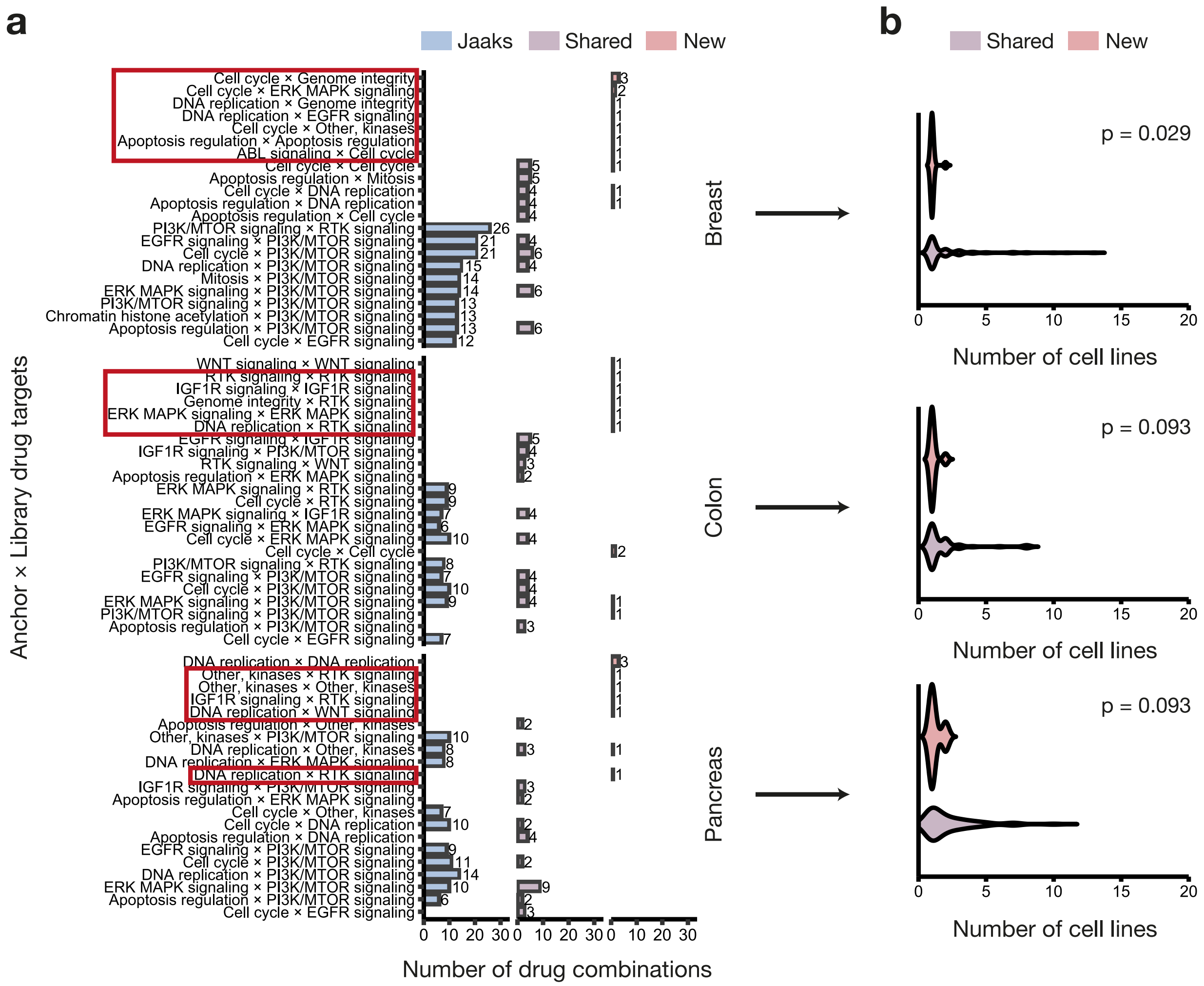


**Supplementary Figure 11. Tissue-specific pathway-pair composition and context-specificity of statistically significant drug combinations.** (**a**) Bar plots show the top 10 most frequent co-targeted pathway pairs among statistically significant drug combinations (ZIP ≥ 10, $p \leq 0.01$). Pathway pairs were defined using canonical ordering of anchor and library targets (e.g., Pathway A × Pathway B) to avoid duplicate representations. For each tissue and combination set, the number of unique drug combinations targeting a given pathway pair is shown. Only distinct drug pairs were counted, and combinations with missing pathway annotations were excluded. Red boxes indicate representative anchor-library target pairs identified exclusively by our statistical framework. (**b**) Violin plots show the distribution of the number of cell lines in which each statistically significant drug combination was identified as synergistic, stratified by tissue and combination set. For each drug pair, the number of sensitive cell lines was counted within the respective tissue. *Shared* combinations denote overlaps with previously reported hits, whereas *New* combinations represent interactions uniquely identified by our statistical framework. The distributions reflect the degree of context specificity across tissues. Statistical differences between *Shared* and *New* combinations were assessed using a two-sided Wilcoxon rank-sum test within each tissue, with Benjamini-Hochberg correction applied across tissues. Here, *Shared* denotes combinations significant under our framework that overlap with the Jaaks et al. hits, whereas *New* denotes combinations significant only under our framework; context specificity is evaluated using our significance definition.

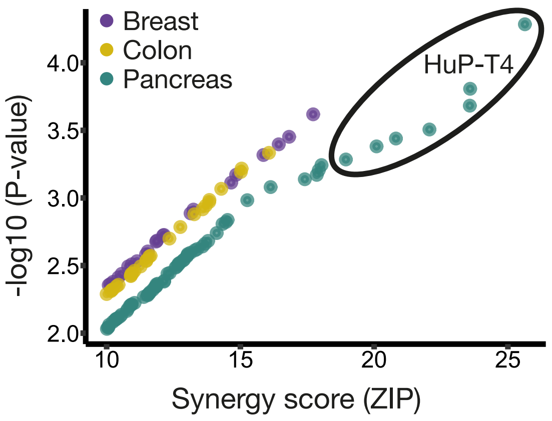


**Supplementary Figure 12. Cross-tissue enrichment of ERK/MAPK-targeting synergistic drug combinations.** Scatterplots summarize statistically significant synergistic observations involving drug combinations targeting components of the ERK/MAPK signaling pathway across breast, colorectal, and pancreatic cancer cell lines. Each point represents a significant drug-pair observation in an individual cell line, stratified by tissue of origin (Breast *n* = 22, Colon *n* = 27, Pancreas *n* = 65; total of 54 unique ERK-targeting drug pairs across tissues). The HuP-T4 pancreatic cancer cell line is highlighted, as it consistently exhibited the strongest synergy among ERK/MAPK-targeting combinations in the pancreatic dataset.

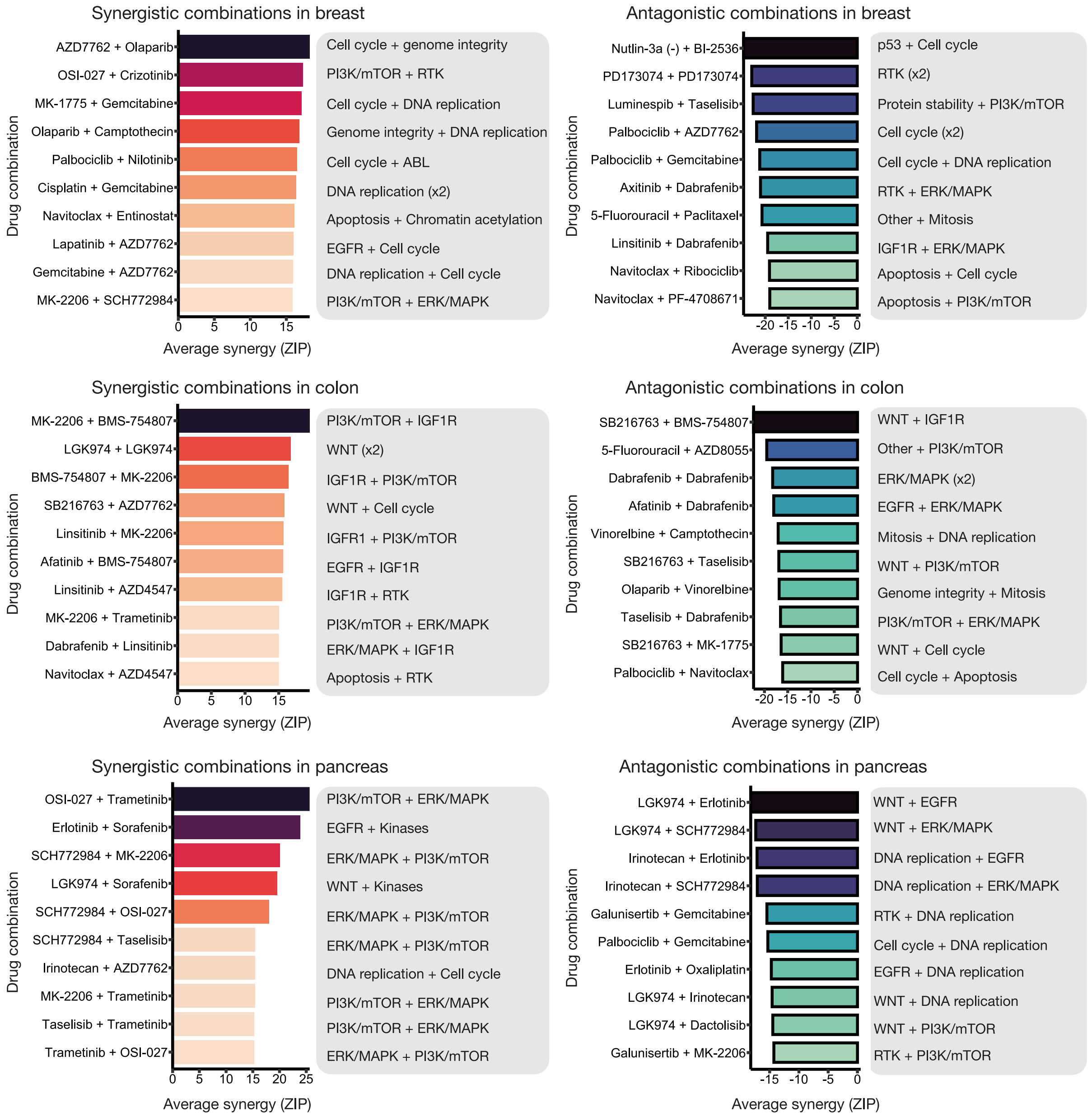


**Supplementary Figure 13. Top-ranked synergistic and antagonistic drug combinations across cancer tissues.** The top 10 synergistic and antagonistic drug combinations identified in breast, colorectal, and pancreatic cancer cell lines from the Jaaks et al. dataset are shown, ranked by their average ZIP synergy scores across all tested cell lines (x-axis). Color shading reflects the magnitude of synergy or antagonism, while pathway labels indicate the biological targets of both anchor and library drugs.

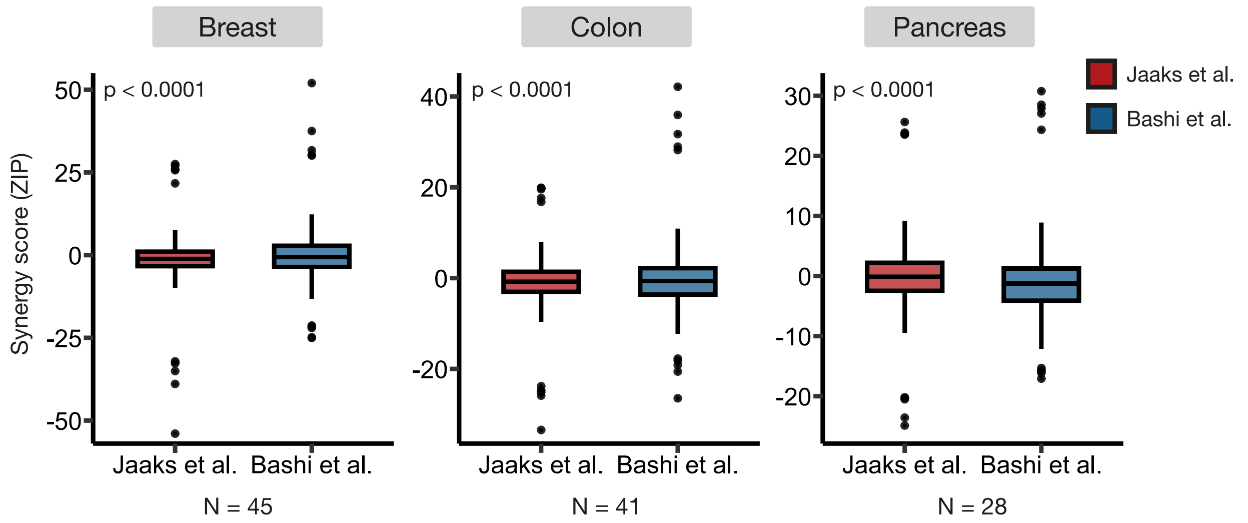


**Supplementary Figure 14. Comparison of synergy score distributions across cell lines shared between datasets.** Boxplots show the distribution of synergy scores across breast, colorectal, and pancreatic cancer cell lines shared between the Jaaks et al. and Bashi et al. datasets. Boxes indicate the interquartile range (IQR) with medians shown as thick horizontal lines, while extreme synergistic and antagonistic combinations are highlighted as individual points. Statistical differences between distributions were assessed using the two-sample Kolmogorov–Smirnov test. *N* denotes the number of shared cell lines per tissue.

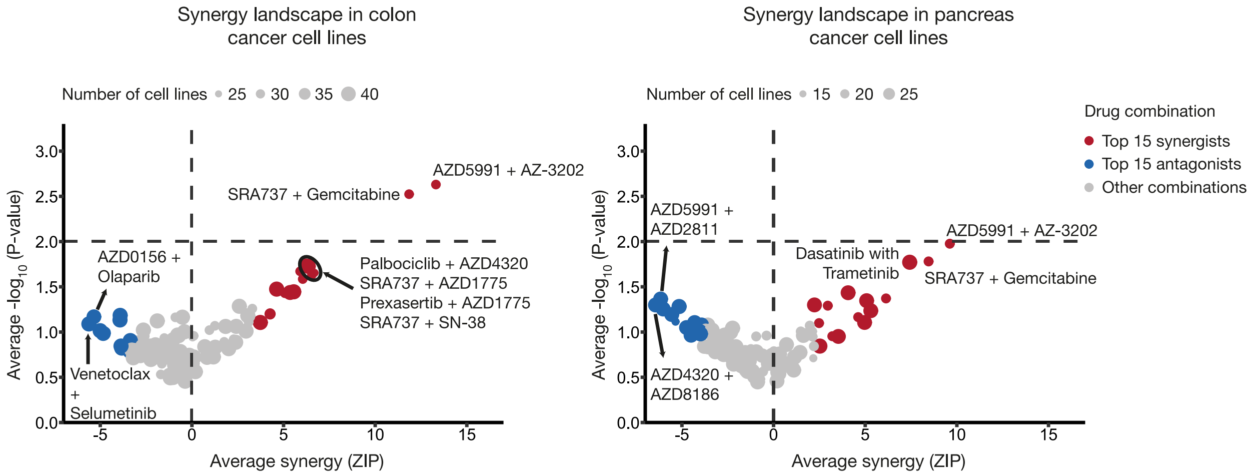


**Supplementary Figure 15. Volcano plots of average ZIP scores against the average *p*-values in the colon and pancreatic cancer cell lines from the Bashi et al. dataset.** The top 10 synergistic and antagonistic drug combinations identified in breast, colorectal, and pancreatic cancer cell lines from the Jaaks et al. dataset are shown, ranked by their average ZIP synergy scores across all tested cell lines (x-axis). Color shading reflects the magnitude of synergy or antagonism, and pathway labels denote the biological targets of both anchor and library drugs.

### Supplementary Tables

**Supplementary Table 1. Pairwise Pearson correlations between synergy metrics across cancer tissues.** Pearson correlation coefficients (r) were computed between synergy metrics across all cancer cell lines from breast, colorectal, and pancreatic tissues. Correlations were calculated using complete cases for each pairwise comparison.

| Tissue | ZIP vs Bliss | ZIP vs HSA | ZIP vs Loewe | Bliss vs HSA | Bliss vs Loewe | HSA vs Loewe |
|---|---|---|---|---|---|---|
| Breast | r = 0.99 | r = 0.80 | r = 0.24 | r = 0.81 | r = 0.25 | r = 0.34 |
| Colon | r = 0.99 | r = 0.70 | r = 0.21 | r = 0.71 | r = 0.22 | r = 0.43 |
| Pancreas | r = 0.99 | r = 0.83 | r = 0.29 | r = 0.84 | r = 0.29 | r = 0.36 |

**Supplementary Table 2. Enrichment of synergistic and antagonistic drug combinations across datasets and tissues.** The proportion of synergistic (ZIP ≥ 10) and antagonistic (ZIP ≤ −10) drug combinations in the Jaaks et al. and Bashi et al. datasets are reported for each tissue type. Enrichment was assessed using Fisher's exact test, with odds ratios indicating the direction and magnitude of enrichment and *p*-values denoting statistical significance.

| Tissue | Synergistic (Jaaks) | Non-synergistic (Jaaks) | Synergistic (Bashi) | Non-synergistic (Bashi) | Odds ratio | *P*-value |
|---|---|---|---|---|---|---|
| Breast | 282 (0.45%) | 62040 | 177 (4.39%) | 3856 | 10.10 | $1.97 \times 10^{-93}$ (<0.0001) |
| Colon | 144 (0.51%) | 27885 | 168 (4.11%) | 3916 | 8.31 | $1.78 \times 10^{-68}$ (<0.0001) |
| Pancreas | 181 (0.94%) | 19071 | 78 (2.89%) | 2620 | 3.14 | $1.74 \times 10^{-14}$ (<0.0001) |

| Tissue | Antagonistic (Jaaks) | Non-antagonistic (Jaaks) | Antagonistic (Bashi) | Non-antagonistic (Bashi) | Odds ratio | *P*-value |
|---|---|---|---|---|---|---|
| Breast | 868 (1.39%) | 61454 | 109 (2.70%) | 3924 | 1.97 | $1.26 \times 10^{-09}$ (<0.0001) |
| Colon | 458 (1.63%) | 27571 | 118 (2.89%) | 3966 | 1.79 | $1.31 \times 10^{-07}$ (<0.0001) |
| Pancreas | 221 (1.15%) | 19031 | 88 (3.26%) | 2610 | 2.90 | $1.51 \times 10^{-14}$ (<0.0001) |

**Supplementary Table 3. Summary statistics of ZIP synergy score distributions across datasets and tissues.** Descriptive statistics of ZIP synergy scores are reported for breast, colorectal, and pancreatic cancer cell lines in the Jaaks et al. and Bashi et al. datasets. *N* denotes the total number of cell lines, *n* the total number of drug combination tests, SD the standard deviation, and IQR the interquartile range.

| Tissue | Screen | Min | Mean | Median | Max | SD | IQR |
|---|---|---|---|---|---|---|---|
| Breast | Jaaks<br>(n=51, n=62322) | -53.96 | -1.13 | -1.11 | 27.51 | 3.73 | -3.29-1.08 |
| | Bashi<br>(n=45, n=4033) | -25.02 | -0.14 | -0.58 | 52.00 | 5.73 | -3.58-2.83 |
| Colon | Jaaks<br>(n=45, n=28029) | -33.44 | -0.82 | -0.82 | 19.98 | 3.80 | -3.05-1.36 |
| | Bashi<br>(n=43, n=4084) | -26.47 | -0.44 | -0.69 | 42.14 | 5.46 | -3.69-2.12 |
| Pancreas | Jaaks<br>(n=29, n=19252) | -24.88 | -0.12 | -0.11 | 25.64 | 3.94 | -2.44-2.22 |
| | Bashi<br>(n=29, n=2698) | -17.08 | -1.04 | -1.23 | 30.77 | 5.12 | -4.05-1.3 |

### Supplementary Files

**Supplementary File 1:** High-throughput screening plate quality control statistics for all screened plates, including associated anchor drugs, library drugs, and tested combinations.

**Supplementary File 2:** Empirical *p*-values and false discovery rate adjusted significance of drug combinations with target annotations.

**Supplementary File 3:** ZIP synergy score matrices and associated drug-cell line annotations across cancer tissues**.**

**Supplementary File 4:** Tissue-specific ranked drug combinations with average ZIP synergy scores and target annotations.

**Supplementary File 5:** Frequency summaries of synergistic and antagonistic anchor and library drug combinations across tissues.

**Supplementary File 6:** Overlap of synergistic drug combinations between the empirical *p*-value framework and the Jaaks et al. synergy detection method.

**Supplementary File 7:** Summary of synergistic and antagonistic drug combinations, anchors, libraries, and cell lines across tissues.

**Supplementary File 8:** Pathway-level aggregation of synergistic drug combinations with average ZIP scores and frequencies across tissues.

**Supplementary File 9:** Reference-based empirical *p*-values and ZIP synergy scores for drug combinations in the Bashi et al. dataset.
